# Correspondence analysis for dimension reduction, batch integration, and visualization of single-cell RNA-seq data

**DOI:** 10.1101/2021.11.24.469874

**Authors:** Lauren L. Hsu, Aedín C. Culhane

## Abstract

Effective dimension reduction is essential for single cell RNA-seq (scRNAseq) analysis. Principal component analysis (PCA) is widely used, but requires continuous, normally-distributed data; therefore, it is often coupled with log-transformation in scRNAseq applications, which can distort the data and obscure meaningful variation. We describe correspondence analysis (CA), a count-based alternative to PCA. CA is based on decomposition of a chi-squared residual matrix, avoiding distortive log-transformation. To address overdispersion and high sparsity in scRNAseq data, we propose five adaptations of CA, which are fast, scalable, and outperform standard CA and glmPCA, to compute cell embeddings with more performant or comparable clustering accuracy in 8 out of 9 datasets. In particular, we find that CA with Freeman-Tukey residuals (CA-FT) performs especially well across diverse datasets. Other advantages of the CA framework include visualization of associations between genes and cell populations in a “CA biplot,” and extension to multi-table analysis; we introduce *corralm* for integrative multi-table dimension reduction of scRNAseq data. We implement CA for scRNAseq data in *corral*, an R/Bioconductor package which interfaces directly with single cell classes in Bioconductor. Switching from PCA to CA is achieved through a simple pipeline substitution and improves dimension reduction of scRNAseq datasets.

## Introduction

Single cell mRNA sequencing (scRNAseq) simultaneously measures the transcript levels of genes in thousands of individual cells, providing a window into the transcriptional and functional diversity of cells in a tissue or experiment. These complex datasets are orders of magnitude larger than those encountered when analyzing “bulk” RNAseq data from tissue samples. While such fine resolution data have the potential to reveal new biological findings, scRNAseq data exhibit sparsity, noisiness, and technical artifacts beyond those seen for bulk RNA samples ^1,2^, necessitating scRNAseq specific pre-processing and normalization^3,4^. Typically scRNAseq analysis includes the use of dimension reduction to attenuate noise and ensure computational tractability, but the choice of method considerably influences downstream analyses, results, and conclusions^3,5^.

Selecting an appropriate dimension reduction method is important; an effective method finds a representation of the data that minimizes noise and redundancy, while uncovering meaningful signals that reveal latent structures and patterns within the data^6,7^. When defined from scRNAseq data, reduced dimension embedding representations are most useful when they preserve meaningful, biologically relevant variation; are robust, meaning that the decomposition of new but similar observations consistently yields a similar embedding space; and generalize and transfer to new data, enabling new observations arising from similar biological processes to be projected into the same latent space.

ScRNAseq counts are generally modeled as multinomially distributed, and are often approximated as negative binomial or Poisson^2^, reflecting the fact that the data are neither continuous nor approximately Gaussian. As such, use of principal component analysis (PCA) requires that discrete and sparse scRNAseq count data be transformed prior to dimension reduction with this method^6^. PCA is a linear dimension reduction method that obtains a low-dimensional data representation along orthogonal linear axes such that the proportion of variance accounted on each axis is maximized in Euclidean space^4,8–11^. Because PCA is most suitable for continuous data that is approximately normally distributed, it may exhibit artifacts when applied to data with gradients or non-continuous data (such as counts); one such artifact, called the “arch” or “horseshoe” effect, occurs when PCA is applied to scRNAseq data without log-transformation^4,6,12^. So, in practice, and despite known issues with applying log-transformation to scRNAseq count data^2,13,14^, most single cell workflows begin with a log(x+1) transformation of the counts matrix, and then use PCA to decompose the resulting “logcounts” data^3^. The use of logcounts has poor theoretical justification, and in some cases may obscure meaningful variation^2,14^, but the resulting reduced dimension embeddings of the data from PCA are nonetheless used in scRNAseq clustering, trajectory analysis, and cell type classification^3^. Several dimension reduction approaches tailored for scRNAseq counts have been proposed, including methods like ZINB-WaVE, the first method appropriate for use with counts which is based on a zero-inflated negative binomial model for decomposition of counts, and zero-inflated factor analysis (ZIFA)^2,15–17^. Still, PCA remains the most widely used method largely due to its simplicity, speed, and computational efficiency. In a comparison of 18 dimension reduction methods, PCA ranked highly when accuracy and performance in downstream analysis were considered with computational scalability^18^.

Classical matrix factorization methods, including PCA, are instances of the general duality diagram approach proposed by Benzécri and the French school of multivariate statistics in the 1970’s^8,19–23^, which pivots focus from the matrix as columns of fixed variables to the matrix as an operator between inner product spaces, unifying classical multivariate methods like PCA with modern kernel methods into the same framework^8,21^. Another matrix factorization method that emerges in the duality diagram framework is correspondence analysis (CA), a fast dimension reduction method appropriate for non-negative, count-based data and can identify relationships between categorical data types that is popular among ecologists for analyzing species-by-site abundance count matrices^8,24^. In practice, PCA is often computed by singular value decomposition (SVD) of column-centered or Z-score normalized data (Figure 1A)^4,25^ and CA is computed by SVD of the Pearson residuals to reveal the row-column associations that deviate from expectation^26^. The principal components in CA partition the co-dependence between the rows and columns such that a higher weight indicates a stronger dependence or association between row and column; for scRNAseq data, CA principal components can identify co-dependence between gene expression counts and particular cells. From this perspective, the main difference is the space into which the data are transformed then decomposed. Whereas PCA partitions the variance in Euclidean space, CA partitions the total contingency chi-square table along linear additive components^27^. CA has a long tradition in diverse settings and disciplines, including linguistics, business and marketing research, and archaeology^26,28^, where it is applied to and further optimized for large, sparse count data. CA has also been applied in bioinformatics to perform codon usage analysis^29,30^; to analyze microarray transcriptomics data^31^; to integrate GO labels with microarray data^32^; and to analyze metagenomic and microbiome data^33^. In *made4*, Culhane et al. implemented CA for microarray and bulk RNA-seq data^34–36^. We now propose its application to scRNAseq analysis.

**Fig 1.**
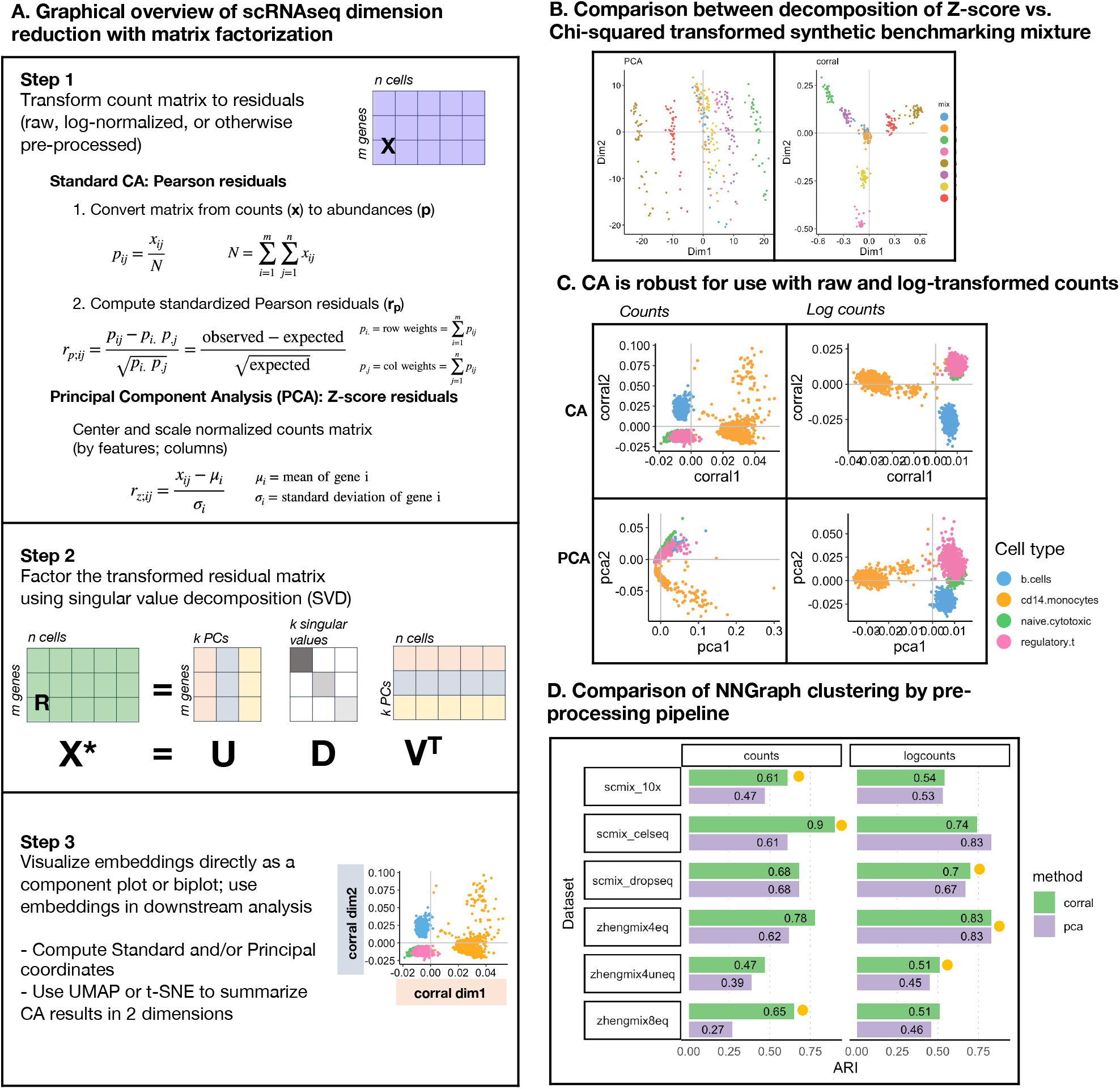
Correspondence Analysis (CA) is a count-based alternative to PCA that is robust for use with raw and log-normalized counts. Correspondence Analysis (CA) is an alternative to PCA for count data that is robust for use with raw and log-normalized counts. **A**. Graphical overview of steps for dimension reduction with matrix factorization, including standard CA and PCA. Standard CA and PCA can be computed with singular value decomposition (SVD) of the Pearson or the Z-score residuals, respectively. **B.** Plots show the first two components generated from PCA (on logcounts; left) and from CA (*corral* on counts; right) applied to a synthetic benchmarking mRNA mixture with 8 groups (data distributed in the CellBench R package; adapted from ^3^). “Cells” are colored by group. CA resolves the groups into clusters, whereas standard PCA is driven by a gradient in the second component and fails to resolve the groups. **C.** Plots show the first two components generated by CA (*corral;* top row) and PCA (bottom row) on both counts (left column) and logcounts (right column) of the Zhengmix4eq dataset, which comprises approximately 4,000 purified PBMCs in approximately equal mixtures. Cells are colored by type. CA is robust for use with counts or logcounts, whereas PCA on counts results in a horseshoe (arch) effect. **D.** CA (green) and PCA (purple) were applied to counts (left column) and logcounts (right column) from six benchmarking datasets (SCMixology; Zhengmix). Embeddings from all approaches were used as input for NNGraph clustering, with performance in recovering published clusters assessed using Adjusted Rand Index (ARI). CA consistently meets or exceeds performance of PCA. Orange circles mark highest ARI achieved in each dataset.

Focusing on the issues of log-transforming scRNAseq counts when applying PCA, Townes et al. (2019), Hafemeister & Satija (2019), and Lause et al. (2021) presented approaches to scRNAseq analysis based on Pearson residual normalization as an alternative to distortive log-transformation. Townes et al. (2019) proposed glmPCA, a generalization of PCA that minimizes deviance rather than mean squared error (MSE) and accommodates non-canonical link functions, and that can be approximated with PCA of Pearson or deviance residuals^2^. Lause et al. proposed analytic Pearson residual normalization^14^, extending work from Hafemeister & Satija, who used a regression-based approach to computing Pearson residuals^13^. Lause et al. cited our open-source Bioconductor workshops which describe CA; the relationships among CA, PCA, and SVD; and their application in scRNAseq data as support that glmPCA from Townes et al., (2019), SCTransform from Hafemeister & Satija (2019) and their approach are CA or closely approximate CA^14,37^. However, CA, which can be computed by SVD on the standardized Pearson residuals, may not be the most appropriate approach when there is overdispersion in the contingency table^38^.

We propose and evaluate five adaptations of CA to address overdispersion in scRNAseq counts. We benchmark the performance of each of these compared to standard CA and with glmPCA^2^, a popular method in the field. In particular, we find that CA with Freeman-Tukey residuals, an alternative chi-squared statistic, is especially performant across a variety of test cases. Because cell clustering and characterization is a key part of most scRNAseq workflows, we set as the goal of the benchmarking task to find embedding representations that facilitate identifying and annotating complex populations of cells. We show that the CA biplot provides a geometric interpretation of features and objects in the same space, which in turn facilitates efficient exploratory data analysis and cluster interpretation. We implemented standard and adapted CA for scRNAseq in *corral*, an R/Bioconductor package that interfaces directly with Bioconductor classes (including *SingleCellExperiment*). Designed for computational scalability, *corral* is fast and performant compared to PCA and other dimension reduction methods, including glmPCA. Switching from PCA to CA with *corral* is achieved through a simple pipeline substitution and improves dimension reduction of scRNAseq datasets.

## Results

### Correspondence analysis: Count-based dimension reduction

Standard correspondence analysis (CA) casts scRNAseq read counts in a contingency table analysis framework and in its canonical form can be conceptualized as a two-step procedure (graphically outlined in Figure 1A; detailed in **Methods**). The count matrix is first transformed to Pearson chi-squared residuals, and the resulting residual matrix is then factored with singular value decomposition (SVD).

CA analysis of scRNAseq does not require, but is compatible with, log-transformed read counts (logcounts). PCA, which has been widely used, requires data transformation, and is therefore generally applied to logcounts data, even though log-transformation of scRNAseq counts distorts latent space representation such that the first dimension is driven by individual cell sparsity, or the number of features with zero observed counts (“zero fraction”)^2^. Since we propose CA as a more suitable alternative to PCA for finding cell embeddings, we compared CA to the widely used correlation-based PCA^4^.

We applied both CA and PCA to a ground-truth scRNAseq benchmarking data set (on both counts and logcounts) obtained by CEL-seq2 sequencing of pseudo-cell mixtures comprising mRNA from eight distinct groups^39^. Figure 1B shows the first two principal components for both PCA and CA. The first PCA component clearly separated cells from three of eight clusters, but PC2 only captures a gradient within the groups. In contrast, CA clearly clustered and separated all groups within two components. Similarly, results in purified PBMCs (Zhengmix4eq benchmarking dataset) demonstrated that CA can be applied directly to counts or to logcounts and still achieve good clustering and separation, whereas PCA on counts produces an “arch” or “horseshoe” effect, arising from the presence of a latent sequential ordering or gradient^12,25^. PCA on logcounts performed similarly to CA on either counts or logcounts.

CA is robust when applied to either counts or logcounts data, obviating the need for log-transformation and avoiding its associated issues. We compared the performance of the four pipeline configurations presented in Figure 1C (CA and PCA on counts and logcounts) on six reference benchmark datasets— three scRNAseq datasets from SCMixology (known cell mixture of three cancer lines sequenced with three technologies)^39^ and three Zhengmix PBMC datasets^40,41^. (Datasets listed in the Benchmarking section of **Methods**.) Cluster recovery based on the annotated cell types in the study was assessed using Adjusted Rand Index (ARI), which assesses similarity between two sets of data partitions (Figure 1D). In all comparisons, CA outperforms or matches PCA’s performance (orange circle indicates highest ARI per dataset).

### Comparison of CA approaches that address overdispersion

CA can be influenced by “rare objects” or outliers^38^. Due to high underlying heterogeneity of gene expression within and between various cell types, scRNAseq data often include biologically “real” outliers as opposed to artifacts due to noisy data. For example, professional secretory cells have a distinct biological profile often driven by extraordinarily high production of one or two proteins, such as insulin in pancreatic islet cells or immunoglobin in immune cells. Similarly, senescent or quiescent cells differ in gene expression profile compared to rapidly dividing cells or high-grade tumor cells.

We propose and evaluate five unique adaptations of CA to address overdispersion in scRNAseq counts. In total, six CA methods (standard CA and the five adaptations) were applied to nine datasets, including the three Zhengmix human PBMC benchmarking datasets, as well as cells from human pancreas, human brain, and *Xenopus* tail (Table 1). Cluster recovery performance on cell embedding representations generated from each specific method was compared and benchmarked in reference to glmPCA^2^, based on the partition similarity of the new clusters with the original annotated cell populations from each dataset (measured with ARI; detailed in **Methods** – Benchmarking).

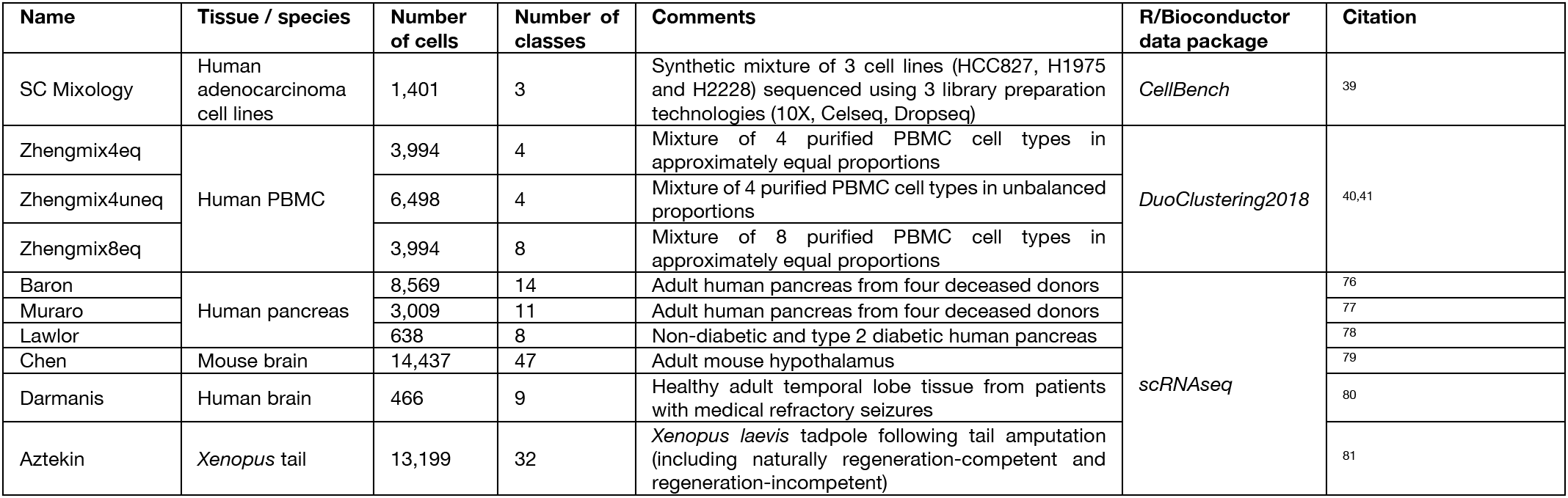

| Name | Tissue / species | Number of cells | Number of classes | Comments | R/Bioconductor data package | Citation |
| --- | --- | --- | --- | --- | --- | --- |
| SC Mixology | Human adenocarcinoma cell lines | 1,401 | 3 | Synthetic mixture of 3 cell lines (HCC827, H1975 and H2228) sequenced using 3 library preparation technologies (10X, Celseq, Dropseq) | *CellBench* | 39 |
| Zhengmix4eq | Human PBMC | 3,994 | 4 | Mixture of 4 purified PBMC cell types in approximately equal proportions | *DuoClustering2018* | 40,41 |
| Zhengmix4uneq |  | 6,498 | 4 | Mixture of 4 purified PBMC cell types in unbalanced proportions |  |  |
| Zhengmix8eq |  | 3,994 | 8 | Mixture of 8 purified PBMC cell types in approximately equal proportions |  |  |
| Baron | Human pancreas | 8,569 | 14 | Adult human pancreas from four deceased donors | *scRNAseq* | 76 |
| Muraro |  | 3,009 | 11 | Adult human pancreas from four deceased donors |  | 77 |
| Lawlor |  | 638 | 8 | Non-diabetic and type 2 diabetic human pancreas |  | 78 |
| Chen | Mouse brain | 14,437 | 47 | Adult mouse hypothalamus |  | 79 |
| Darmanis | Human brain | 466 | 9 | Healthy adult temporal lobe tissue from patients with medical refractory seizures |  | 80 |
| Aztekin | *Xenopus* tail | 13,199 | 32 | *Xenopus laevis* tadpole following tail amputation (including naturally regeneration-competent and regeneration-incompetent) |  | 81 |

The five adaptations of CA fall into three general approaches (Figure 2A). The first class of approaches was to explicitly apply a variance-stabilizing transformation to the count matrix prior to computing Pearson residuals. Lause et al. (2021) discussed variance-stabilizing transformation as compared to Pearson residual normalization, though in their study did not combine variance stabilization and Pearson residual normalization prior to matrix decomposition. They reported that the degree of correction from variance-stabilizing transformation alone was insufficient for scRNAseq data in their pipeline configuration and found that only normalizing with analytic Pearson residuals was more effective than only applying variance stabilization^14^. Given that scRNA-seq counts are often approximated as Poisson-distributed, we considered three variance-stabilizing transformations that are typically applied to count data. These three square-root based transformations all originate from R.A. Fisher’s observation that performing an arccosine transformation on the square root of multinomial probabilities yields approximately normally distributed angles on a hypersphere^42^. The first was square root transformation of count data (Row 3 of Figure 2A), which has been used to correct overdispersion in Poisson counts^43^. The second is Anscombe’s variance-stabilizing count transformation (Row 4 of Figure 2A), originally proposed in 1948 for use with Poisson, binomial, and negative binomial data^44^. Third, we used the Freeman-Tukey variance-stabilizing count transformation (Row 5 of Figure 2A), originally proposed in 1950, also for Poisson and other count data^45^.

**Fig 2.**
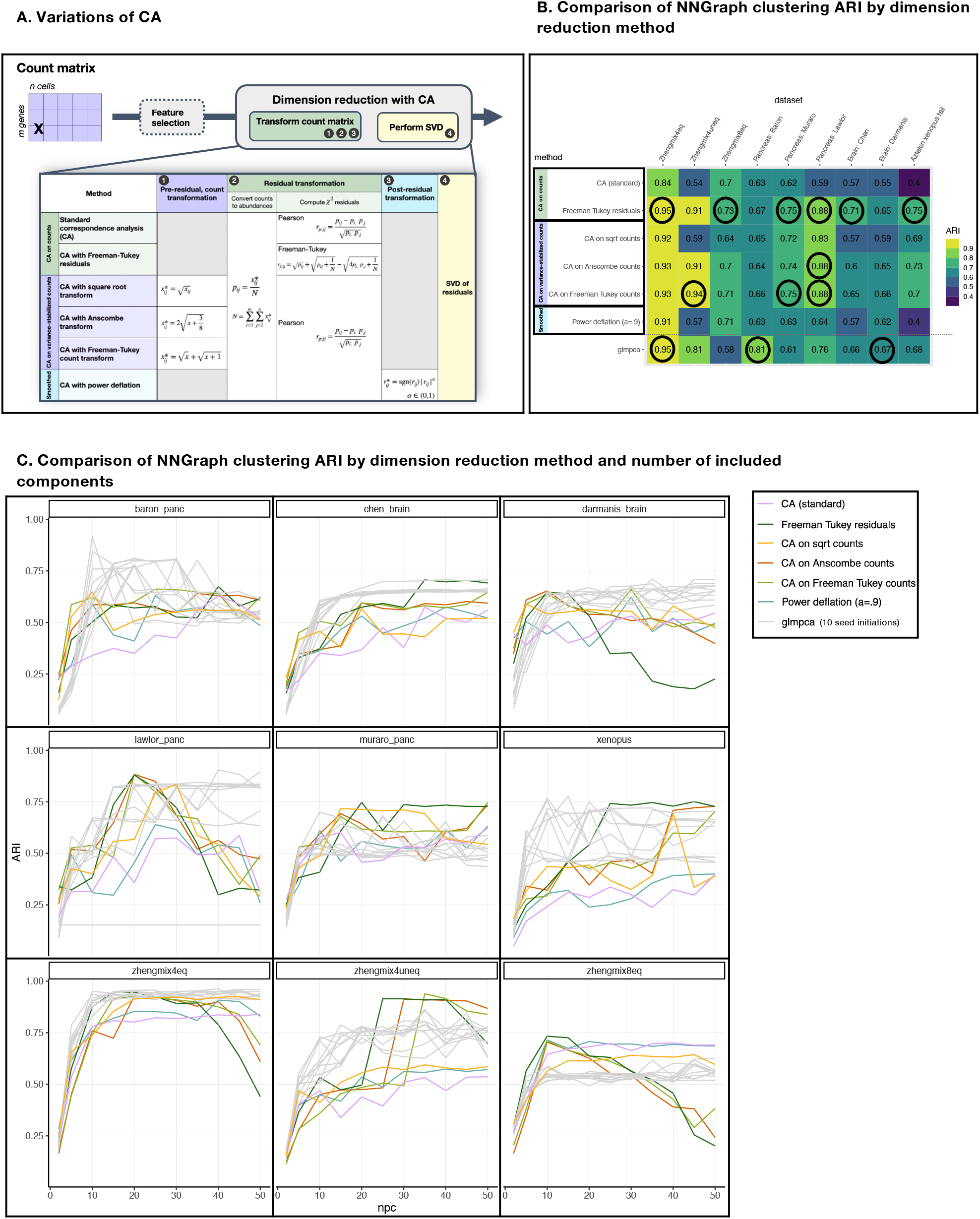
CA can be adapted to address overdispersion in count data. CA adaptations to address overdispersion in count data. **A.** Table summarizing the standard CA procedure and five adaptations to address overdispersion. The first set (row 1 and 2) include methods that involve no transformations apart from computing chi-squared residuals. The second set (rows 3-5) feature variance-stabilizing transformations performed on counts prior to standard CA. The third approach (row 6) smooths the chi-squared residual matrix with a minor “power deflation” prior to decomposition with SVD. **B.** Table of NNGraph cluster recovery performance achieved by each method (rows), in nine datasets (columns), reporting the maximum ARI selected across a range of PCs (full results of ARI by PC shown in Figure 2**C**), with ARI from ten runs of glmPCA were averaged prior to selecting the maximum. Highest ARI (to two decimal places) in each dataset is circled, and the cell clusters in the original datasets are used as the reference groupings. Freeman-Tukey residuals exhibit the best overall performance, with the highest ARI in 6 of the 9 datasets. **C.** Plot of ARI by number of components in each of nine datasets (same as **B**), colored by method. Results for glmPCA (gray) include ten seeds.

Our results indicate that variance stabilization improves performance of standard (classical) CA. Variance stabilization of counts prior to computing Pearson residuals provided great gains in downstream clustering with ARI increases of 0.4 in two studies (Zhengmix4uneq, Aztekin *Xenopus* tail); square-root transformation prior to CA increases ARI in 7 datasets, while transformation to Anscombe counts or Freeman-Tukey counts increased ARI in every dataset when compared to standard CA (with no variance stabilization of counts prior to computing Pearson residuals). Indeed, Anscombe’s variance-stabilizing count transformation achieves the highest observed ARI in 1 of 9 test datasets (pancreas: Lawlor) and Freeman-Tukey variance-stabilizing count transformation had best overall performance in 3 of 9 datasets (Zhengmix4uneq; pancreas: Muraro, Lawlor). Although the square root count transformation did not outperform the other two transformations in any of the comparisons, its ARI was within 0.05 of other two transformation in 7 of 9 datasets. Furthermore, in the pancreas datasets, variance-stabilizing count transformation coupled with standard CA yielded the highest ARI overall, outperforming glmPCA.

The second variation we considered is “power deflation” as a data smoothing method. Power deflation handles extreme outliers in the chi-squared residual matrix by raising all transformed residual values to a power, a, prior to performing SVD, while preserving sign (Bottom row of Figure 2A). Conceptually, this procedure is similar to the Tukey ladder transformation^46^, and has a smoothing effect on the matrix of chi-squared distances, reducing the impact of outlying values while preserving the ordering of values. To achieve a “soft” smoothing effect, we considered *α* ∈ [0.9, 0.98] (data not shown) and present results for *α* = 0.9 in Figure 2. This approach is similar to the classic square root variance stabilizing transformation for Poisson counts, with the special case where *α* = 0.5, but it differs in that the transformation is applied to the chi-squared residual matrix after rather than to the count matrix. In all nine datasets, this power deflation smoothing approach performed comparably to, or better than, standard CA, although its impact on CA performance was less than variance-stabilizing count transformation.

Third, we considered an alternative chi-squared statistic that is better-suited to count data with high levels of sparsity and overdispersion. CA with Freeman-Tukey residuals (CA-FT) has been applied to archaeological site data, where it exhibited a variance-stabilizing effect and outperformed standard CA (SVD of the Pearson residuals), in the analysis of sparse, over-dispersed artifact data (counts of archeological artifacts by site)^45,47,48^. Both Pearson residuals and Freeman-Tukey residuals are members of the Cressie-Read family of power divergence statistics for testing goodness-of-fit in multinomially-distributed count data, and when squared, both residuals are chi-square distributed random variables^47,49^. We found that CA-FT is well-suited for scRNAseq counts (Row 2 of Figure 2A), outperforming standard CA in all nine datasets and its performance was comparable to (ARI within 0.02) or superior to glmPCA in 8 out of 9 benchmarking datasets. In most datasets CA-FT also had higher or comparable clustering accuracy (ARI) to standard CA with variance-stabilizing transformation. CA-FT achieved the highest ARI overall in 6 out of 9 datasets. Unlike standard CA, we observed little benefit to combining CA-FT with variance-stabilizing transformation (square root, Anscombe, or Freeman-Tukey) (Figure S1); while the performance of standard CA improves dramatically with variance-stabilizing transformation, CA-FT adjusts for and is appropriate to be used with overdispersed data.

Component selection can greatly influence downstream cell clustering analysis, so we considered clustering performance as a function of the number of components selected (Figures 2C, S2). The ability to recover “known” clusters (measured with ARI between clustering output and the published cell types) was higher for the simpler mixtures of known, purified cell types (Zhengmix datasets). For the complex tissues examined (Brain; Pancreas; *Xenopus* tail), the “true” number of cell types are experimentally estimated from the scRNAseq data. There was heterogeneity in the number of cell types described in the same tissue between different studies, possibly because cell annotations can be assigned at low resolution (e.g., T-cells), or at high resolution (e.g., CD4 T-cells, exhausted CD8 T-cells, etc.), depending on the particular study question. For instance, the pancreas datasets Lawlor, Muraro, and Baron described eight, eleven, and fourteen cell types in their respective analyses (Table 1). We observed an association between the number of components and the complexity of the clustering task. More components may capture more total variation in data and thus might increase performance when performing higher resolution annotation. Figure 2C shows that more components generally increased ARI in more complex tissue. However, for datasets where the reference cell type annotations are lower resolution (fewer cell types), including more components could reduce the ARI since their results will be higher resolution (more cell types) and therefore technically less concordant with the original reference. This reveals a limitation of current benchmarking approaches. A new method could find biologically meaningful groups, but perform poorly if scored using ARI on low resolution benchmarking datasets. We observed in our results that the Lawlor and Darmanis datasets, both annotated at lower resolution, showed the steepest decline in ARI clustering performance when more PCs are included.

In contrast, there was little gain and, for some, a reduction in ARI with more components in the Zhengmix datasets, which comprise combinations of distinct PBMC cell types sorted and purified prior to sequencing. In simple datasets, including additional components beyond those that sufficiently capture the biological variance may add stochastic, technical, or systematic noise in the system. Benchmarking each of the methods with ranking by maximum ARI was robust to the number of components; CA-FT was consistently most performant, whether the first thirty or fifty (Figures S2,2B) components were included in downstream clustering.

CA, CA-FT, and other variations generate a nearly deterministic result that is stably reproduced. In contrast, glmPCA is not deterministic, and therefore results may vary substantially when the method is rerun on the same dataset (Figures 2C,S3). For reproducibility, we tested ten random seed initiations of glmPCA (Figure 2C), which revealed that glmPCA results are consistent for simpler datasets but in other datasets, such as the *Xenopus* tail dataset, performance varies dramatically between iterations. In the Lawlor pancreas dataset, one iteration failed, suggesting that results were somewhat dependent on finding a “lucky seed.” In simpler datasets, such Zhengmix, all methods generated high ARI scores and glmPCA results had consistency between individual runs (Figure 2C). However, there was greater variation in glmPCA performance with increasing data complexity. For each dataset, we present the average of the maximum ARI achieved in each of 10 runs of glmPCA.

CA variations adapted for overdispersion outperform standard CA or glmPCA in downstream clustering (Figure 2B). Of the approaches we considered, CA-FT was most performant, outperforming standard CA with variance-stabilizing transformation and the power deflation approach.

### Geometric interpretation of cell and feature embeddings

The CA biplot provides a natural framework for cluster interpretation, highlighting biologically meaningful relationships among gene expression patterns and cell populations, and may be extended to guide feature selection. Every transformed count (residual) in a CA matrix has an intuitive interpretation, as it is the chi-squared test statistic for strength of association between a particular row (expression of a gene) and column (cell). The CA matrix captures the strongest associations between gene expression and cells, highlighting functional contrasts by individual cells and by subpopulations of cells. Biplots visualize associations between features and objects, or in this case, genes and cells. Rather than examining the feature and object embeddings individually, the biplot places both sets of embeddings on the same axes, revealing both the associations that may exist among either rows or columns separately, and also between particular rows and columns^6,50^. Distance from the origin indicates the magnitude of association; the angular rotation distance (cosine similarity) reflects similarity of the cells (or genes) to each other, or association between cells and genes.

We performed standard CA on the Zhengmix8 PBMC benchmarking dataset, plotting the first two dimensions of the resulting cell and gene embeddings (Figure 3). The 20 genes with highest weight by L2 norm in the first two dimensions are colored blue, with a corresponding gene label. Cell populations are colored by cell type. The biplot highlights genes that have strong associations with and may discriminate between particular cell populations. For example, natural killer (NK) cells constitutively express granulysin, encoded by the gene GNLY, and although they are not exclusive producers of granulysin, GNLY expression in other cells, like cytotoxic T-cell populations, is driven by immune activation^51^. The CA biplot shows that GNLY has a high weight in PC2 (far from origin) and has a similar angular rotation as the NK cell population (high cosine similarity). Correspondingly, the inset ridge plots in Figure 3 showing histograms of log expression in cell populations confirm it is highly expressed specifically in the NK cell population.

**Fig 3.**
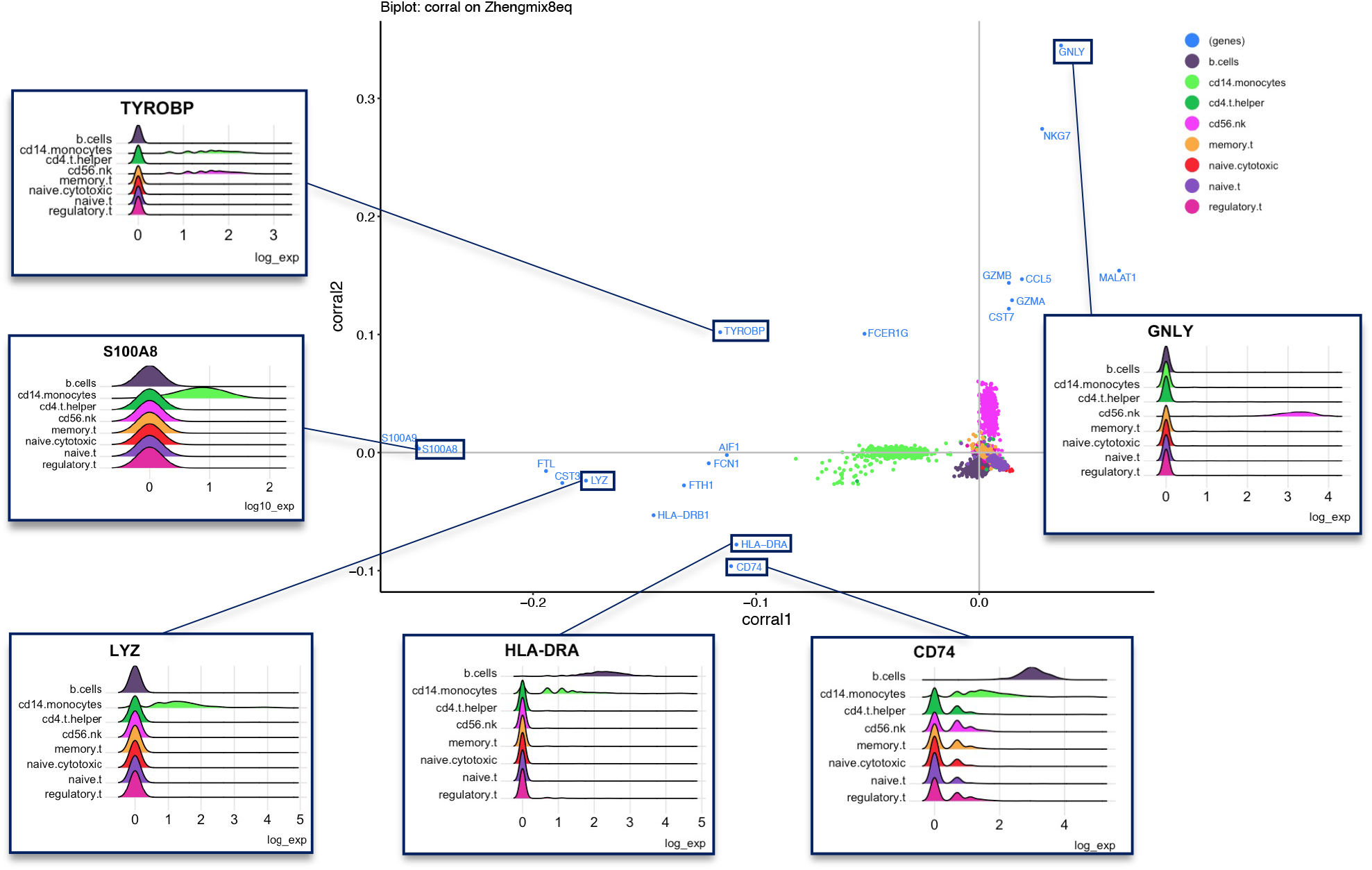
Geometric interpretation of correspondence analysis: Illustrating associations between genes and cell populations. Geometric interpretation of correspondence analysis: Illustrating associations between genes and cell populations. Biplot of the first two dimensions of CA in the Zhengmix8 dataset. The eight cell populations are colored by type, while genes are labeled and colored in blue. The top twenty genes by weight (furthest from the origin in the first two components) are shown. Six biologically significant genes are highlighted, and ridge plots illustrate their log-expression: GNLY is highly expressed in NK cells, whereas TYROBP is highly expressed in both NK and CD14 monocytes. LYZ and S100A8 are both highly expressed, monocyte-specific genes. Both CD74 and HLA-DRA are highly expressed in B cells, and moderately expressed in monocytes, as shown in the respective ridgeplots.

Calcium-binding proteins S100A8 and S100A9 (MRP8 and MRP14 respectively) are constitutively expressed in monocytes and neutrophils^52,53^. Correspondingly, in the CA biplot in Figure 3, the expression of both genes is strongly associated with the monocyte population (same direction, large magnitude), consistent with the relative log-expression of S100A8 among cell populations (inset plot). Similarly, LYZ encodes for lysozyme, a molecule highly secreted by monocytes^54^. Reflecting the elevated differential expression of the gene among the monocyte population shown in the inset, the gene is far from the origin while also close in angle to the cell population.

Biplots also inform about genes highly and differentially expressed in multiple cell populations: TYROBP encodes for a signaling adaptor protein (KARAP/DAP12), which was initially identified as a wiring component in NK anti-viral and anti-tumoral function^55^. TREM-1, a KARAP/DAP12-associated surface protein, amplifies monocyte, macrophage, and granulocyte activation by cytokines and chemokines following LPS stimulation^55^. While other lymphoid and myeloid cells may express TYROBP, it has predominantly been observed in NK, monocytes/macrophages, and dendritic cells, consistent with the enriched expression levels in the expected cell types: NK and monocytes. The gene is projected between these cell populations; expression ridge plots confirm that it exhibits elevated expression specifically in NK and monocyte cell populations.

CD74 is part of the MHC class II complex, consistent with both its biplot positioning and expression plot: angularly, it lies closest to the B cell population, but is also rotated slightly towards the monocyte population^56^. Correspondingly, expression of CD74 is seen in cells of all types but is most elevated in B cells and in some monocytes. Similarly, HLA-DRA encodes the alpha chain of the HLA-DR protein, which is a cell-surface receptor in the MHC class II complex^57^. Both B cells and monocytes are professional antigen presenting cells that require all the machinery of the MHC class II complex, so these genes are important for function of both cell types, and both genes in the biplot are angled between the most relevant cell types, providing a biologically meaningful summary of associations between genes and cell sub-populations.

The CA biplot facilitates unified analysis of cell and gene embeddings, which can inform cluster interpretation and serve as a basis for integrating with (and extending) other methods, such as gene set enrichment analysis and projection of supplementary data into a shared latent space.

### Integrative multi-dataset dimension reduction with corralm

The need to integrate cells from multiple batches motivates continued refinement and development of CA^10,35,58^. Our multi-table adaptation of CA, implemented as *corralm* in the *corral* R/Bioconductor package, operates using indexed or Freeman-Tukey residuals, and finds a joint multi-table embedding. It is suited for light to moderate integration tasks (e.g., different sequencing runs of an experiment). For complex integration tasks with substantial batch effects, corralm may not fully integrate the data because it is a multi-table extension of CA dimension reduction, and is not optimized for batch integration and contains no explicit integration step. Since CA embeddings can be easily substituted for PCA in a pipeline, we investigated whether inclusion corralm in batch integration improved the performance of popular integration methods that include a PCA step. For example, widely used batch correction methods, FastMNN and Harmony, include a PCA step. We compared corralm’s performance with widely used batch integration methods (Figure 4), including LIGER^59^, MNNCorrect, Harmony, and Seurat (suggested pipeline including SCTransform normalization and CCA integration), all of which performed well in recent benchmarking studies^59–63^. To assess corralm as a PCA pipeline substitute, we included in the comparisons corralm coupled with Harmony and MNN.

**Fig 4.**
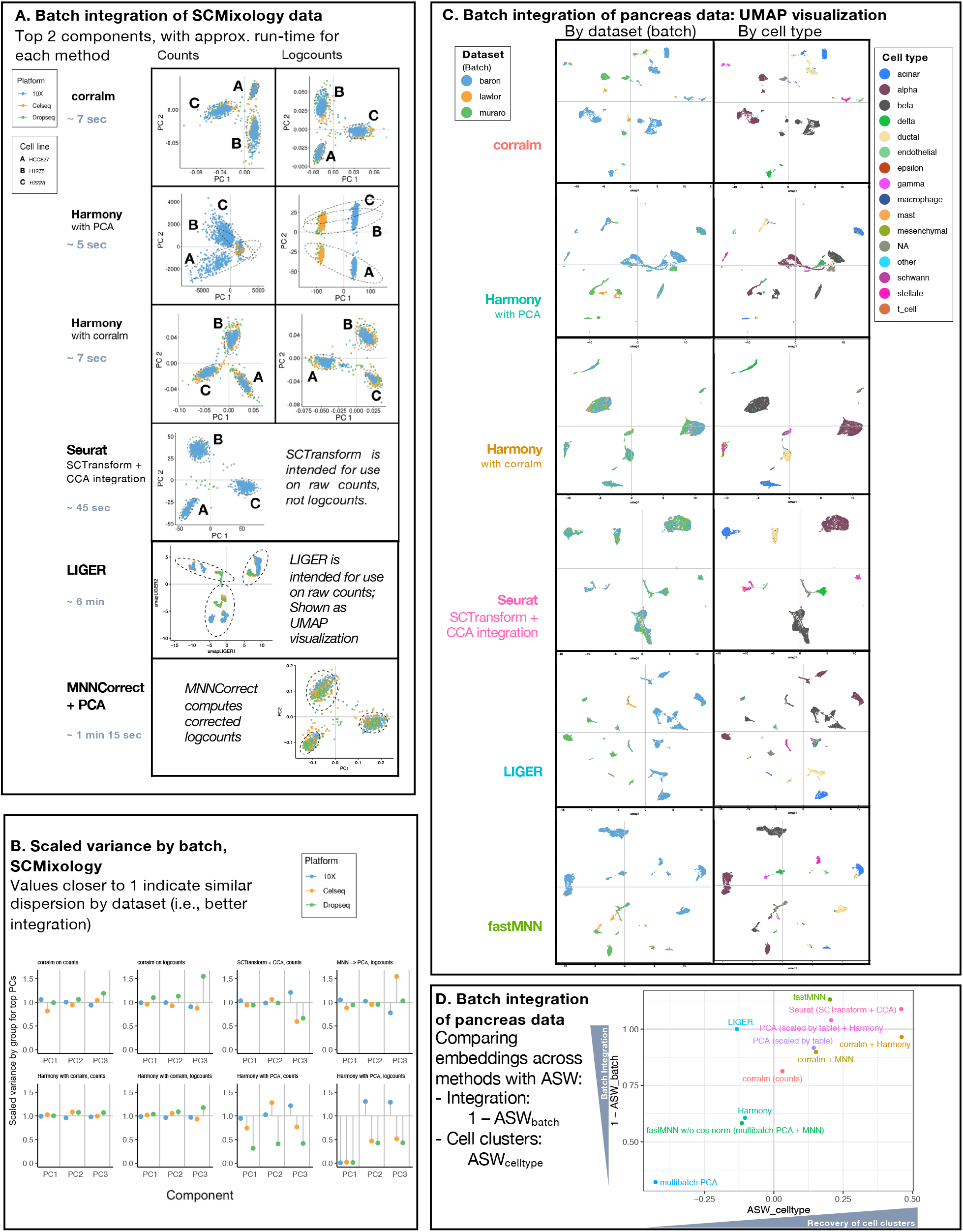
corralm, a multi-table adaptation of CA, finds joint embeddings across batches by decomposing into a shared, low-dimensional latent space. The corralm multi-table adaptation of CA integrates count matrices across batches by finding a shared, low-dimensional latent space. **A.** Comparison of nine integration workflows on the SCMixology benchmarking dataset (comprising mixtures of three cell lines: H2228, H1975, and HCC827 that were each used with three library preparation protocols—Dropseq, Celseq2, and 10X—followed by Illumina sequencing) The first column shows results on counts, and the second column shows logcounts (where appropriate). corralm is both fast and performant and can be combined with methods such as Harmony (the 3^rd^ row) to further improve performance. **B**. Scaled variance (SV) of the batches representing the three SCMixology library preparation platforms, computed on the first three components of counts and logcounts presented in Figure 4A, colored by batch. SV close to 1 indicate that embeddings exhibit similar distribution across batches. corralm, Harmony with corralm, and SCTransform exhibit good batch alignment, while Harmony with PCA shows values far from 1, suggesting that the embeddings were not successfully integrated across batches (Includes all methods with ranked components). **C.** Batch integration of pancreas data. For each of a selected set of methods, the left column shows UMAPs colored by dataset (batch), while the right column shows UMAPs colored by cell type. **D.** ASW_cell type_ assesses the embedding based on preserving biological context, while 1 - ASW_batch_ assess integration, and are on the x and y axes respectively. For all methods, this is computed on 8 PCs.

First, to compare performance in a clear and simple ground-truth scenario, each method was applied to batch integration of the SCMixology benchmarking dataset comprising scRNAseq profiles from a mixture of three cell lines (H2228; H1975; HCC827), obtained in three batches using different library preparation platforms (Dropseq; Celseq2; 10X)^39^. Second, to compare performance in a more complex, biologically realistic example, the methods were applied to integration of three human pancreas datasets, obtained on different platforms in separate studies: Baron, Lawlor, and Muraro (detailed in **Methods** – Benchmarking below).

In the SCMixology dataset, the “ground truth” is unambiguous, and we expect the low-dimensional representation to align data across batches and identify distinct cell line clusters. Figure 4A shows the first two components of the reduced dimension representation of results from *corralm*, Harmony with *corralm* embeddings, SCTransform with CCA, and MNNCorrect with PCA successfully integrate batches while preserving cell line clusters (Figure 4A, rows 1,3,4,6). In contrast, Harmony (using PCA embeddings, as published) fails at both data integration and cluster detection on these same data (Figure 4A, row 2). LIGER succeeds in cluster separation but fails in integration, as visualized in the UMAP (Figure 4A, row 5). Qualitatively, SCTransform with CCA exhibits the best alignment by batch and tightest clusters by celltype, but its run-time is an order of magnitude slower than *corralm* and Harmony with *corralm*.SCTransform with CCA runs in 45 seconds, while *corralm* and Harmony with *corralm* run in 7 seconds for the equivalent task, allocated one core of a laptop (**Methods**—Benchmarking). LIGER and MNNCorrect are significantly slower, running in approximately 6 minutes and 1.25 minutes, respectively. Although the SCMixology dataset is relatively small (1,401 cells), at scale, this difference in run-time would significantly impact the overall speed of a pipeline, thus demonstrating an advantage of *corralm* and Harmony with *corralm*.

Cluster evaluation measures like ARI assess whether clusters can be re-identified, but do not directly quantify how well datasets are integrated in their low dimensional embedding representations. We propose a new metric, scaled variance (SV), for assessing batch integration of datasets comprising similar cell populations across batches (Figure 4B; detailed in **Methods**). For each dimension of each embedding, we compute the variance of the subset of observations from each batch and scale by the overall variance in that dimension as a measure of under- or over-dispersion of the subset’s embeddings in that dimension. For example, in the SCMixology benchmarking dataset, biologically identical samples were assayed using three library preparation methods (Dropseq; Celseq2; 10X), with each batch expected to have the same distribution of cells. SV values closer to one indicate better integration (more similarity in dispersion) in a given dimension by batch. Consistent with Figure 4A, the SV plots (Figure 4B) showed that SCTransform had the best integration, with all SV points very close to one. Similarly, *corralm* and Harmony with *corralm* also showed good batch integration, and both outperform Harmony with PCA, which had SV values far from one.

In the more complex and realistic pancreas scRNAseq integration task, the performance of data integration methods were assessed qualitatively by comparing UMAPs (Figure 4C and S5) and quantitatively with ASW cluster metrics^64^ (Figure 4D), as in a previous benchmarking study^62^. Assuming that the given cell type labels from each dataset are ground truth, in an embedding where cell types form compact and perfectly separated clusters, ASW_cell type_ should be close to 1. Batch integration was measured by 1 – ASW_batch_, where values near 1 (ASW_batch_ near 0) indicate integration and less clustering by batch. Corralm is a simple joint dimension reduction that includes neither optimization for batch nor explicit batch integration steps, and therefore is not expected to outperform methods optimized for batch correction. However, we see corralm outperforms multibatch PCA (Figure 4D). Moreover, corralm combines well with integration pipelines: pairing Harmony or MNN correction with corralm embeddings improves the embedding as compared to both corralm alone and to the original pipelines with PCA. In Figure 4D, we report that corralm (with Freeman-Tukey residuals) coupled with Harmony exhibits comparable performance to the Seurat routine in terms of integration and biological cluster separation. Qualitatively, these UMAPs are similar (Figure 4C). In contrast, other methods shown in Figure 4C were less successful in integrating the batches, though they did appear to preserve at least some of the biological structure.

### Computational performance of corral’s CA implementation

The *corral* implementation of CA leverages fast, approximate, partial SVD from the *irlba* R package^65^; even when allocated one core on a laptop (**Methods** — Benchmarking), *corral* runs in under a minute for a dataset of 1,500 features and over 20,000 cells (50 components). Figure 5A shows that for the analogous task, glmPCA takes over an hour, and that across a range of dataset sizes (1,500 features), glmPCA’s run-time increases rapidly with the number of cells, while CA (*corral*) scales much more favorably. As SVD implementations improve, run-time and/or memory use may be further reduced by modularly incorporating these into the *corral* pipeline. Standard CA and the variations we considered are not sparse implementations; computational performance may be further enhanced with adaptations for sparsity. Since CA has similar computational requirements to PCA, replacing PCA with CA is a simple pipeline substitution.

**Fig 5.**
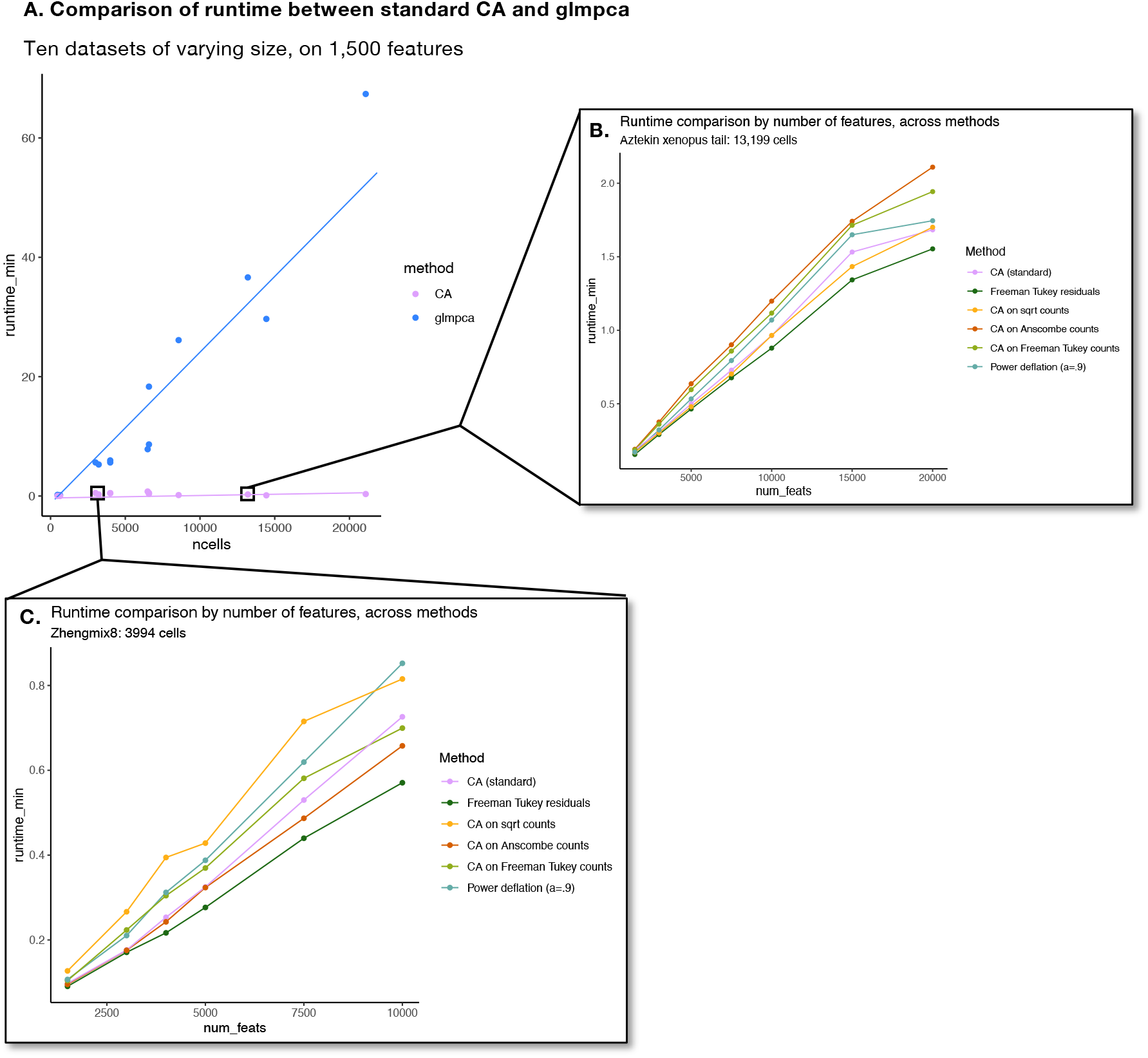
Computational performance of CA and its adaptations. Computational performance of CA and its adaptations. **A**. Plot comparing runtime for standard CA and glmPCA on ten datasets, selecting down to 1,500 features in each. Standard CA consistently runs in under a minute, even for datasets with over 20,000 cells, while glmPCA scales less favorably and requires over an hour for the equivalent input matrix (1,500 features x ~22,000 cells). **B**. Plot comparing runtime with increasing number of features in the Aztekin *Xenopus* tail dataset, across the CA adaptation methods. Since they use similar routines, their runtimes are fairly similar. **C**. Plot comparing runtime with increasing number of features in the Zhengmix8 dataset, across the CA adaptation methods. In both **B** and **C**, it is notable that even with an order of magnitude more features, CA and its adaptations run in a fraction of the time glmPCA takes.

## Discussion

Correspondence analysis (CA) is a statistical technique with a rich theoretical foundation that was first proposed and mathematically characterized nearly a century ago^66^ and which has continued to be developed and extended. CA has been periodically “rediscovered” and adapted in a variety of disciplines^20,28,67–69^ and most recently in the field of scRNAseq analysis: several groups have suggested Pearson residual-based normalization prior to matrix decomposition with PCA^2,13,14^, a routine that is conceptually similar to standard CA—apart from differences in how residuals are computed, one additional distinction in this routine is PCA’s additional Z-score normalization step^4^ after computing Pearson residuals, as opposed to directly decomposing the residual matrix with SVD.

Correspondence analysis with Freeman-Tukey chi-squared residuals (CA-FT) is a simple and effective adaptation of CA for dimension reduction of scRNAseq counts. We compared the performance of CA and five CA variations that address scRNAseq overdispersion, benchmarking these against glmPCA^2^, a popular method in the field. CA-FT was most performant overall in a scRNAseq cluster recovery task. Our analyses also showed that, combined with standard CA (Pearson residuals), incorporating variance-stabilizing transformations and “power deflation” smoothing both improve performance in downstream clustering tasks, as compared to standard CA alone. Therefore, for dimension reduction of scRNAseq data, we recommend using CA-FT or, when using standard CA, incorporating variance-stabilization and/or smoothing.

Data normalization and dimension reduction significantly impact downstream scRNAseq analyses. Performance of dimension reduction approaches depend on variance structure, noise, and other characteristics of a dataset; we find, as has been reported elsewhere^18^, performance of methods vary depending on the characteristics of individual datasets. Benchmarking studies are limited by lack of robust reference datasets reflecting the depth of complexity and nuance in actual biological research; most high-quality, “ground-truth” benchmarking datasets are derived from simple “pseudo”-cell mixtures, or from pools of distinct cell types. Neither reflect the true diversity of cell types in tissues, nor properties of real-world research data. Typically, parameters like number of “true” clusters are unknown *a priori* and depend on the specific research question and context. A complementary approach is to consider benchmarking datasets obtained by sequencing complex tissue samples, although these datasets also have their own disadvantages; cells in such studies are assigned identities based on one analytical method (and for one particular set of study objectives) without a way of independently validating the assignments. Therefore, these single-context annotations set an overly narrow standard for future benchmarking studies of other methods, which can never outperform the method used for initial assignment. With advances in systematic benchmarking frameworks for complex datasets in different contexts, we will be better equipped to test the merits of each approach and identify optimal approaches based on data characteristics.

As such, the analyses we present here are somewhat limited by the context-specific annotations of our benchmarking datasets, since we use as the ground truth labels the original annotations published with these datasets. Except for SCMixology and Zhengmix (both comprising well-defined cell clusters and by design simpler than data from complex tissues), the datasets we analyzed did not have independently validated cell type annotations, so performance is limited by the original cell type assignments. Even if a given method better distinguishes important sub-populations or rare cell types from clustering, these advantages may not be reflected in the ARI, and the method would actually receive a small penalty for differences from “reference.” Given the complexity of and subjectivity inherent in cell cluster annotation, researchers may call different cell populations or clusters from the same dataset, depending upon the research objectives. The diversity of research questions and data challenges in single cell biology necessitate the breadth of statistical and computational approaches. The robust conceptual framework for CA and its empirical performance advantages over PCA argue for its application in scRNAseq analyses.

We implemented CA, CA-FT, and other variations that adjust for overdispersion of scRNAseq data in the R/Bioconductor package *corral* (including documentation, tutorials, vignettes), enabling its integration into commonly used analytical pipelines^3,37^. We conclude with ideas for future development—CA, especially when situated within the broader duality diagram framework, can serve as both a platform for and rich source of further methods development. By simultaneously visualizing both cell and gene embeddings, the CA biplot emphasizes the row-column duality inherent in these data, facilitating joint analysis of genes and cells. The unified approach to analysis of gene and cell embeddings provides a natural framework to extend and/or integrate with other approaches, including gene set enrichment analysis, supervised decomposition, and projection of supplementary data into shared latent space–for example, with a similar approach as used previously in *mogsa* and *om/cade4*^10,34,36^. Embeddings can be used as matrix operators to project supplementary data into shared latent space, enabling multi-modal and multi-batch integration, as well as fast approximation methods. Matrix projection via multiplication is fast and scalable, even for very large datasets, and in future extensions, can serve as the basis for fast, approximate dimension reduction approaches based on decomposing a representative subset of the data and then projecting into the space the full matrix. As advances in library preparation methods enable sequencing of ever-larger numbers of individual cells, computational considerations are critical in selecting analytical methods and designing scRNAseq pipelines.

## Methods

### Standard correspondence analysis on a single table

Similar to many other matrix factorization methods, correspondence analysis comprises two main steps: a data transformation routine (see also Figure 1A), and a matrix decomposition operation (such as SVD or eigen analysis). In applying “standard” CA to scRNAseq count data, we use SVD to decompose Pearson residuals of gene-by-cell expression count matrix, where the residual quantifies the difference between the observed and the expected data. In this case, the expected value is the product of the row and column weight from the original count matrix. A positive residual, indicating that the observed value (count) for that feature/gene and cell pair is higher than expected, suggests an association or codependency; correspondingly, a negative residual shows a lower value than expected, suggesting indicating a negative association between the expression of a gene expression and a cell subpopulation. When squared, the residuals are chi-squared distributed random variables, and their sum of squares comprises a chi-squared goodness-of-fit test statistic with (n-1)(m-1) degrees of freedom^47,70^.

Correspondence analysis is a dual scaling along the rows and the columns of each count matrix.

CA applied to scRNAseq count data proceeds through the following two discrete steps:

1. Transformation from counts to standardized residuals. Suppose X is an *m* × *n* matrix with *n* cells (indexed on *j*) in the columns and *m* features (indexed on *i*) in the rows, comprising observations *X_ij_*. The abundance *p_ij_*, the weight of the *i*th row *p_j_*, and the weight of the *j*th column *p_.j_* for a given observation *x_ij_* are:

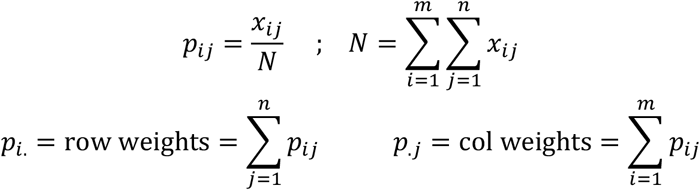 The expected abundance for observation *x_ij_* is p_i_.*p_.j_* and is what we would expect to see in a cell assuming there is no relationship between a row and column. The standardized (Pearson) residuals *r_p;ij_* are the difference between the observed and expected, and can be computed:

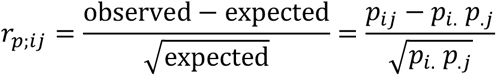 This transformation is equivalent to the computation applied in contingency table analysis of categorical data measuring the strength of association between elements in a row and a column. It yields a matrix M_s_ where the sum of the distances of the points to their centroid (“total inertia”) is the chi-squared statistic of the matrix^26,28^. As a result of this transformation M_s_ is centered and should appear more Gaussian, and therefore is appropriate input for SVD.
2. Matrix decomposition. M_s_ is decomposed using singular value decomposition (SVD) to find left singular matrix **U**, diagonal matrix of singular values **D**, and right singular matrix **V** such that:

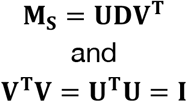 The resulting **U** matrix can be either used as an embedding directly, with each column representing a dimension in the new latent space, or coordinate scores can be computed. Standard coordinate scores are given by dividing the **U** and **V** matrices by the vectors of row weights and column weights, respectively. Principal coordinate scores are given by multiplying the standard coordinate scores by the vector of diagonal values of the matrix **D**. The principal coordinate scores differ from the standard coordinate scores by a scalar on each dimension, and both reflect the ordination scores of the features and cells^38^. Unlike in PCA, where differences in embeddings approximate Euclidean distances, correspondence analysis decomposes the overall chi-squared statistic. The value of the underlying chi-squared statistic is high when there is an association between a row-column pair of the table.

### Variations of CA

We considered five variations of CA to address overdispersion in scRNAseq counts (also summarized graphically in Figure 2A).

1. **CA with Freeman-Tukey chi-squared residuals.** Instead of computing the Pearson residuals described above, the residuals are computed:

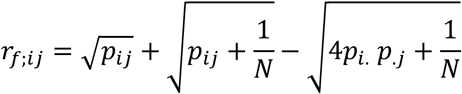 The matrix of these residual values is then decomposed with SVD as described in Step 2 above.
2. **CA with variance-stabilizing transform: Square root.** The square root of the matrix of counts **X** is computed before performing the residual transformation.
3. **CA with variance-stabilizing transform: Anscombe.** Each element *x_ij_* of the matrix of counts **X** is transformed to 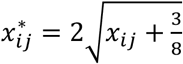. The residual transformation is computed on the variance-stabilized counts matrix **X**\*.
4. **CA with variance-stabilizing transform: Freeman-Tukey.** Each element *x_ij_* of the matrix of counts **X** is transformed to 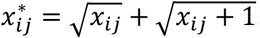. The residual transformation is computed on the variance-stabilized counts matrix **X**\*.
5. **CA with power deflation.** After performing the Pearson residual transformation, each value in the matrix of residuals is transformed to a power of *α* ∈ (0,1), while preserving the sign. Each element *r_ij_* in the residual matrix is transformed to 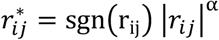. We recommend selecting *α* ∈ [0.9,0.99] for a “soft” smoothing effect, presenting results for *α* = 0.9.

### corralm: multi-table adaptation of correspondence analysis

The adaptation of correspondence analysis for the integration of multiple tables is similar to the method for single tables with additional matrix concatenation operations. When integrating datasets, we employ indexed residuals, by dividing the standardized residuals by the square root of expected proportion to reduce the influence of column with larger masses (library depth), which is a known source of batch effect in scRNAseq studies. Indexed residuals have a straightforward interpretation for example a value of 0.5 indicated that the observed value is 50% higher than the expected value. A value of −0.5 indicated that the observed value is 50% less likely than expected to have a gene-cell association than expected.

1. Match tables and select features. Identify the intersection of features across the *k* matrices to be integrated, and subset the tables for only those *m\** features. While in these analyses we focus on batch integration and therefore match on features, the tables can either be matched by features, for integration across batches, or by cells, for multi-modal integration across ‘omic types.
2. Transformation from counts to indexed residuals. Given each table with *n* cells and *m*\* features, the row weight *p_i._*, column weight *p_.j_*, and abundance *p_ij_* for each observation are computed as described above for standard CA. The indexed residuals *r_ij_* can be computed:

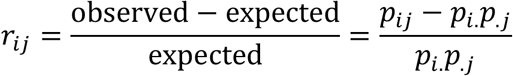 Each table is scaled separately, so as to preserve the internal structure of each dataset.
3. Concatenate matrices. The transformed matrices of indexed residuals are then concatenated along the matching features to form a new matrix **M_c_** which has *m*\* features and the total number of cells in the *k* matrices (i.e., sum of *n* across *k*).
4. Matrix decomposition. Singular value decomposition (SVD) is applied to the concatenated matrix of indexed residuals **M_c_** to find left singular matrix **U**, diagonal matrix of singular values **D**, and right singular matrix **V** such that:

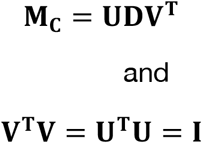 The columns of the **U** matrix then serve as the embeddings generated by this procedure, and the cells correspond to their indices in the concatenated matrix **M_c_**.

Depending upon downstream analysis, it may be important to select an appropriate number of PCs. Similar to PCA, the number of components can be selected using the elbow method with the scree plot, e.g., as implemented in the *findPC* R package (as in Figure 4C for corralm with Harmony)^71^.

### Scaled variance plot

When integrating embedding representations across batches, measures for cluster evaluation are effective for assessing group compactness and recovery of cell populations via clustering. However, they do not directly assess how well dataset embeddings are integrated across batches. To focus specifically on batch integration, we developed and applied a heuristic scaled variance metric, which captures the relative dispersion of each batch with respect to the entire dataset. The scaled variance of component dimension *d*\* for the subset of observations in batch *b*\*, *SV_b*,d_*, is computed with:

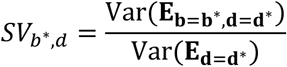

where **E** is the matrix of embeddings, and *b* indexes the rows (observations by batch) while *d* indexes the columns to indicate which component dimension to evaluate. When the datasets are well integrated, SV values for each batch are close to 1, indicating that each batch has similar dispersion as compared to the entire embedding. This metric is appropriate when the types of cells represented in different datasets are expected to be similar but cannot account for situations where the expected distribution of cell types (and therefore, embeddings) is fundamentally different between batches.

### Benchmarking

We considered the ten scRNA-seq benchmarking datasets shown in Table 1. The reduced dimension embeddings from each method were clustered using walktrap nearest neighbor graph clustering, as implemented in the *bluster* package’s default NNGraph parameter set^72,73^. Performance on the clustering task was assessed with Adjusted Rand Index (ARI)^74^, using as “ground truth” the cell type labels from the original datasets. Walktrap was selected as the main method for clustering based on performance; we observed, similar to others, that the walktrap algorithm better preserves hierarchical structure than Louvain clustering and overall achieves higher ARI^75^. Results comparing Louvain clustering and with walktrap clustering are included in Figure S4. We note that whilst some variability in clusters and ARI was observed between runs, CA-FT consistently ranked as the most performant method across the range of datasets. Results shown in Figure 2C are from clustering using different numbers of PCs. Results shown in Figure 2B are computed by taking the maximum across all the tested PCs from Figure 2C, and for glmPCA, the value shown is the average of the maxima achieved by each seed (ten seeds tested in total). Datasets (detailed below) were acquired from three R/Bioconductor data packages: CellBench, DuoClustering2018, and scRNAseq. Links to each of these are included below in the Data Availability section.

Analyses were conducted on a 2019 MacBook Pro laptop (2.3GHz 8-core Intel Core i9; 64GB memory; OS: Catalina). Computational performance is reported based on results from this device.

In the SCMixology integration (Figure 4A, B), each of the benchmarked methods is run with the default settings as suggested in their respective documentation/vignettes. mnnCorrect from the *batchelor* R/Bioconductor package is run on the logcounts matrices, then decomposed with PCA^60^. The LIGER result is shown as UMAP visualization because since it is a NMF-based method, we found that the visualization of the UMAP embeddings directly was challenging since the dimensions of the embedding are not ranked by performance, and are also constrained to only positive values^59^. Similarly, LIGER is not shown in the scaled variance plot for the same reason, and we would not recommend using the scaled variance plot approach with other methods that do not generate ranked components.

In the pancreas integration (Figure 4C, S5), all UMAP plots were generated using *n_neighbors* = 40 or *n_neighbors* = 50. Methods were similarly implemented as in the SCMixology integration results. PCA (scaled by table) was implemented as described in our minireview^4^. Multibatch PCA was performed with the *batchelor* implementation (multibatchPCA), as was the “+ MNN” method (reducedMNN). In the result for corralm + Harmony, the elbow method (implemented in findPC; perpendicular option^71^) was used for PC selection prior to running Harmony^61^. Average silhouette width (ASW) was implemented with the *cluster* R package, using Euclidean distance^64,82^. To enable joint evaluation, labels were harmonized, such that matching cell types are assigned the same label across datasets. In particular, activated stellate and quiescent stellate were merged to stellate; gamma/pp and pp were merged with gamma; duct and ductal were merged.

## Declarations

### Ethics approval and consent to participate

Not applicable.

### Consent for publication

Not applicable.

### Availability of data and materials

Code and documentation are available in the *corral* R/Bioconductor package: https://www.bioconductor.org/packages/corral

R code to reproduce the figures and analysis in this manuscript is available on Github at: https://github.com/laurenhsu1/corral_manuscript

A tutorial describing different implementations of PCA and CA, including *corral*, is available at: https://aedin.github.io/PCAworkshop

The datasets used in these analyses are detailed in the Benchmarking section of **Methods**, including citations and where the data can be accessed directly through R data packages.

For ease of access, links for each Bioconductor data package used in this paper are included below:

- *CellBench:* https://bioconductor.org/packages/release/bioc/html/CellBench.html
- *DuoClustering2018:* https://bioconductor.org/packages/release/data/experiment/html/DuoClustering2018.html
- *scRNAseq:* https://www.bioconductor.org/packages/release/data/experiment/html/scRNAseq.html

### Authors’ contributions

LH and ACC wrote the manuscript and conceptualized the methods presented. ACC wrote the Bioconductor workshop vignette on CA. LH developed the R/Bioconductor package *corral*, wrote code to perform analyses, and created figures.

## Acknowledgements

We are grateful for helpful discussions with Prof. John Quackenbush and his lab at Harvard TH Chan School of Public Health, Prof. Aedín Culhane’s lab at University of Limerick, and with Bioconductor colleagues funded by the Chan Zuckerberg Initiative seed network program. We are also grateful for support from Prof. Judith Agudo and her lab at Dana-Farber Cancer Institute.

