## Supplement for "Correspondence analysis for dimension reduction, batch integration, and visualization of single-cell RNA-seq data"

### Supplementary figures

#### Figure S1

**A.**

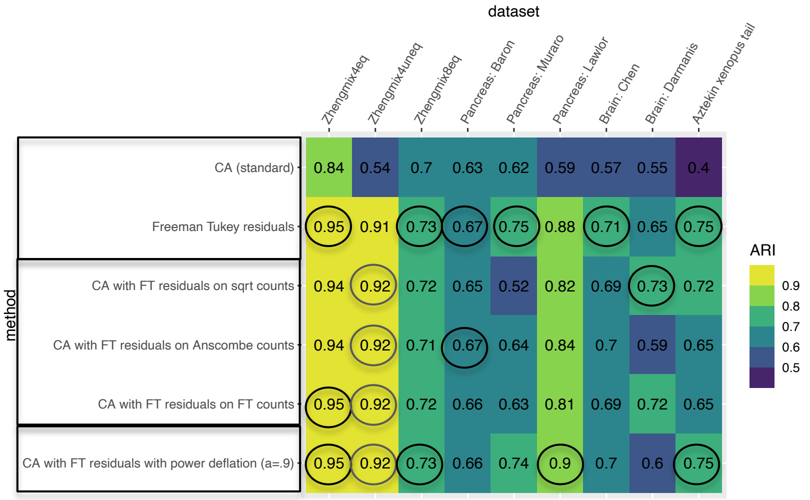


**B.**

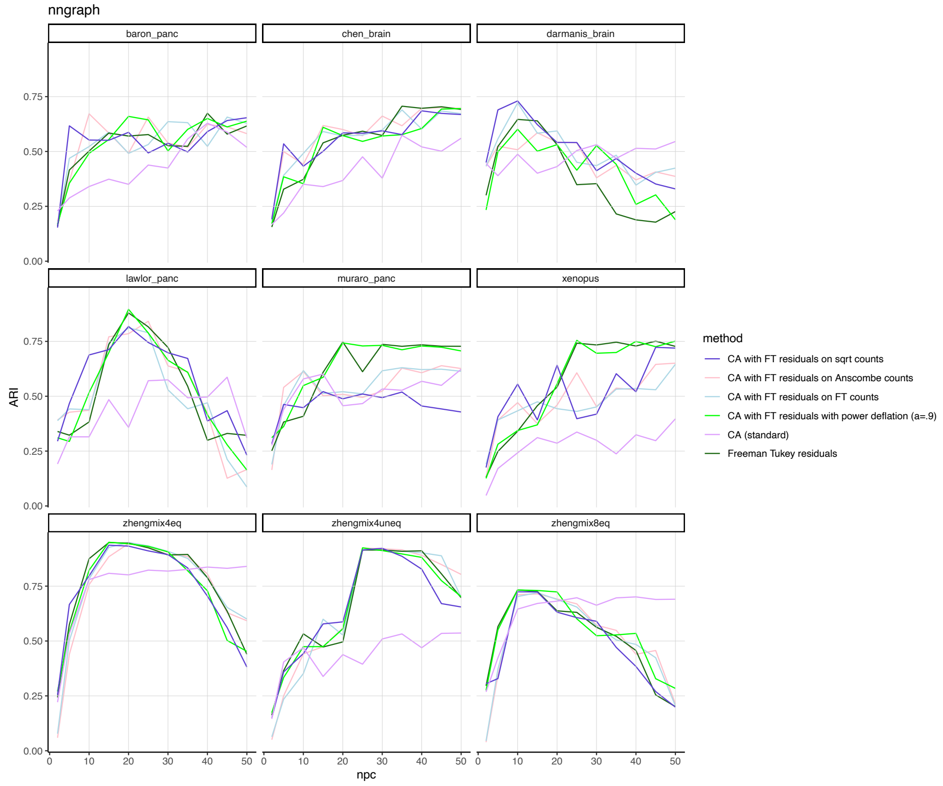


**Figure S1. Correspondence analysis of Freeman-Tukey (CA-FT) residuals with variance stabilization or power deflation** shows limited gains in ARI over CA-FT alone. **A.** Table of NNGraph cluster recovery performance achieved by each method (rows), in nine datasets (columns), reporting the maximum ARI selected across a range of PCs (full results of ARI by PC shown in Figure S1B). Highest ARI (to two decimal places) in each dataset is circled, and the cell clusters in the original datasets are used as the reference groupings. **B.** Plot of ARI by number of components in each of nine datasets (same as **A**), colored by method.

### Figure S2

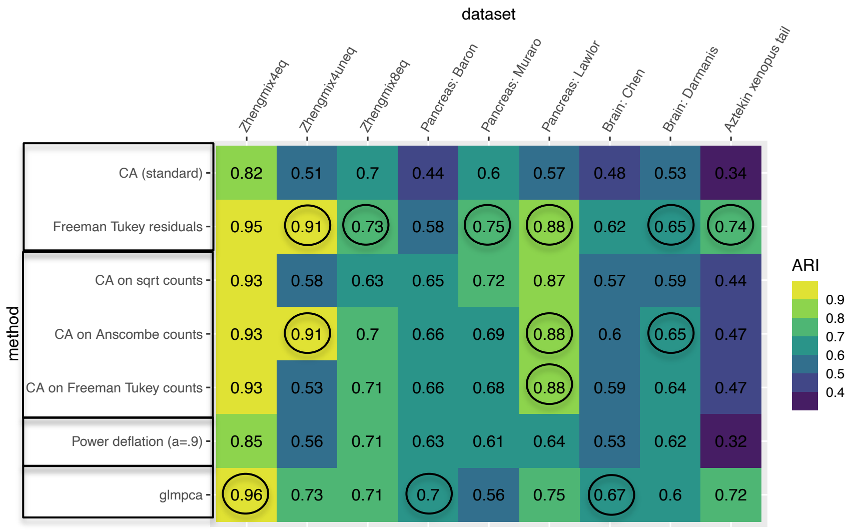


**Figure S2. Testing 30 components.** CA variations are performant with up to 30 PCs (Walktrap). Clustering performance (ARI) of each dimension reduction approach on 9 benchmarking datasets. This figure replicates and is consistent with analysis in Figure 2 but considers 30 PCs (whereas 50 PCs, as included in Figure 2).

#### Figure S3

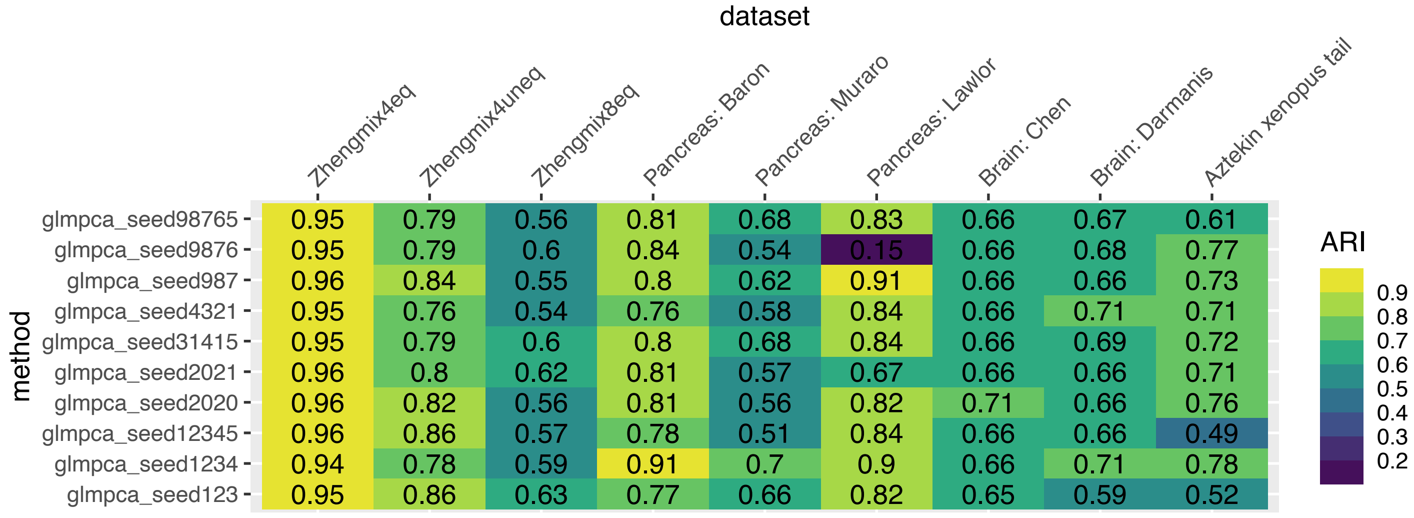


**Figure S3. Stability of glmPCA**. glmPCA of dataset was repeated 10 times with different seed initiation. Heatmap shows the maximum ARI achieved by each glmPCA iteration shown in Figure 2C. The values for glmPCA in Figure 2B are the mean of these values.

### Figure S4

**A.**

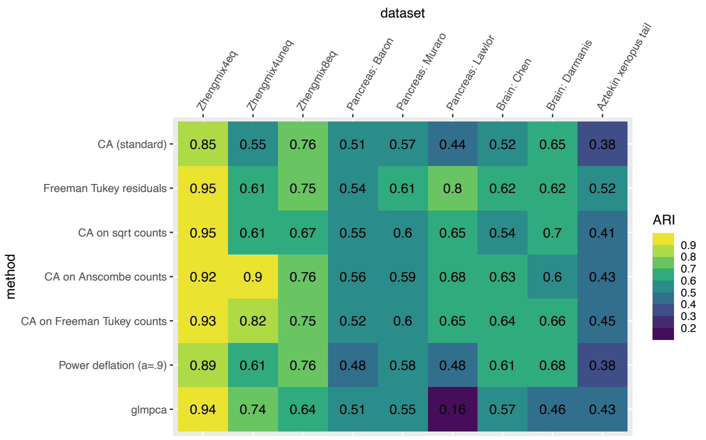


**B.**

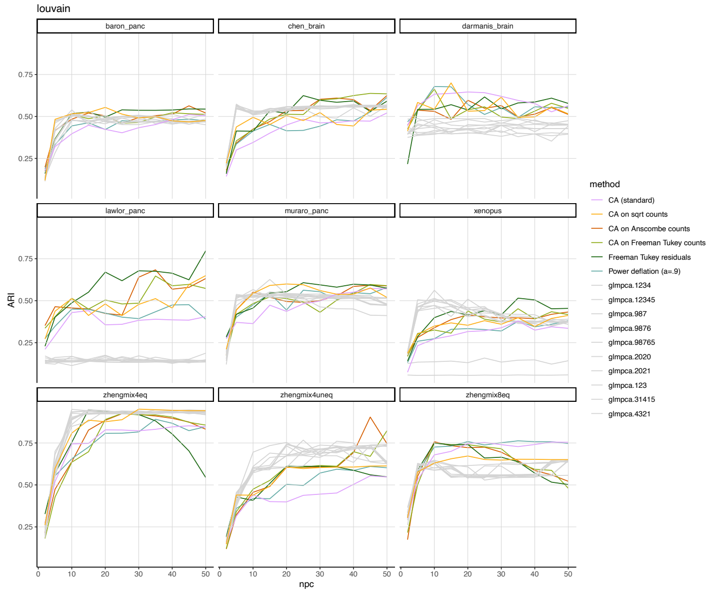


**C.**

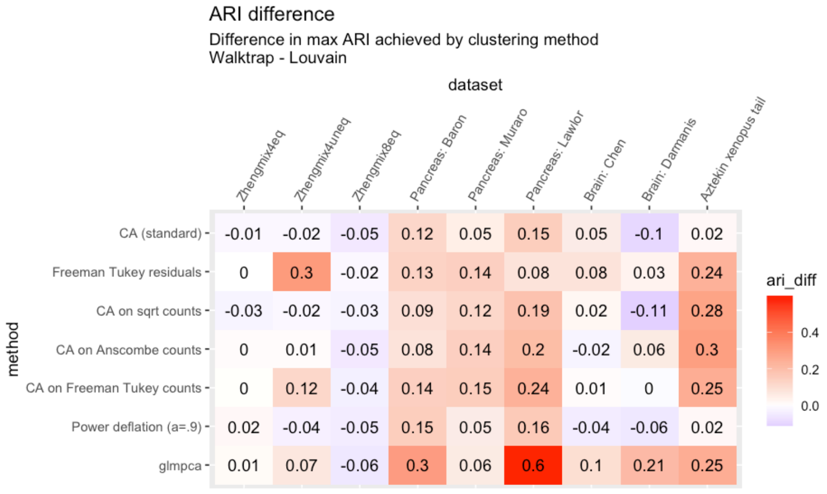

**Figure S4. Comparison of clustering performance between Walktrap and Louvain**. Walktrap showed overall better clustering performance than Louvain, given limited parameter tuning. A. Analogous clustering results as in Figure 2B in main text, but using Louvain instead. B. Analogous clustering result as in Figure 2C in the main text. C. Difference between Figure S4A (max ARI using Walktrap) and Figure 2B (max ARI using Walktrap). Statistically significant difference using signrank test ($p = 1.5e-5$)

#### Figure S5

##### A. corralm (Freeman-Tukey residuals) on counts

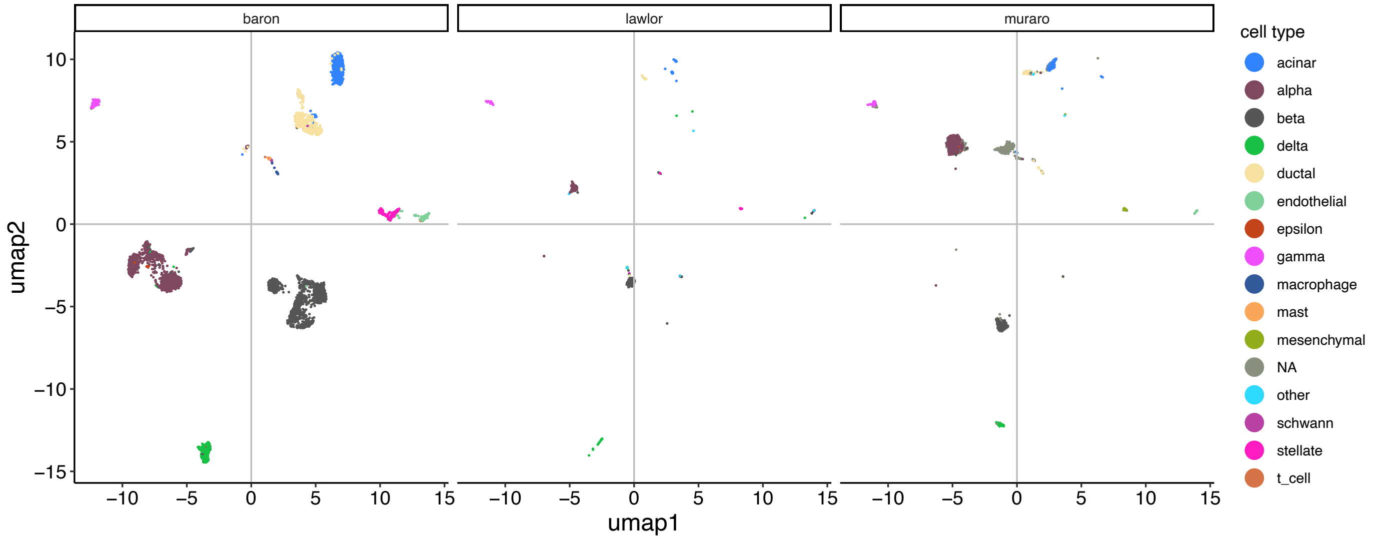


##### B. Harmony with PCA

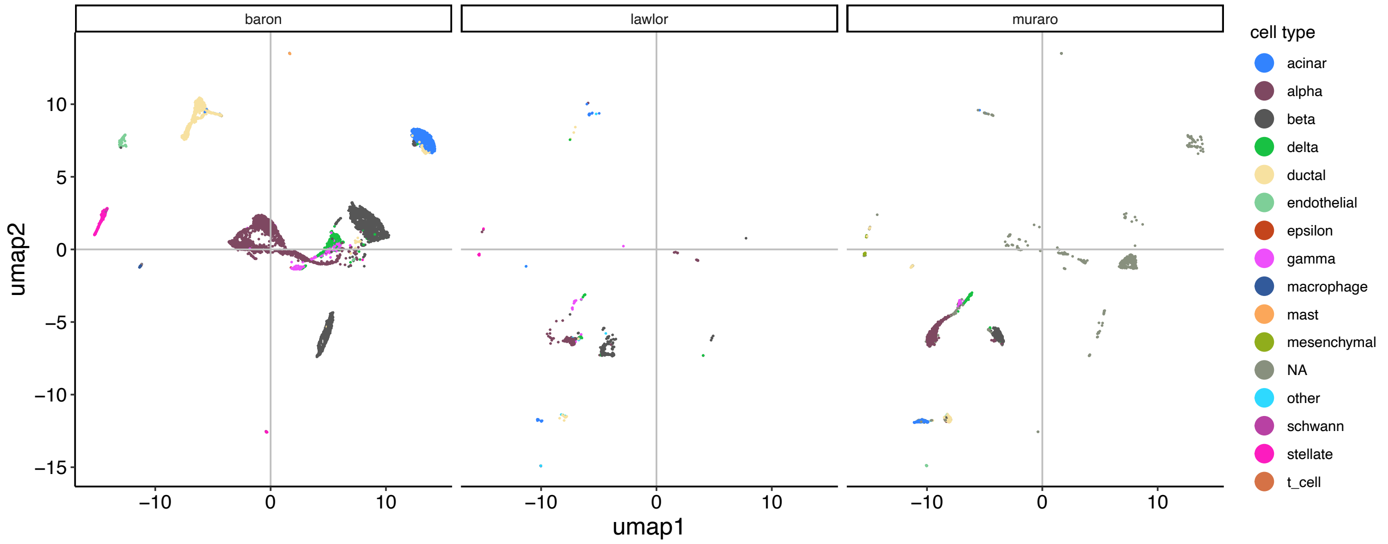


##### C. Harmony with corralm (Freeman-Tukey residuals)

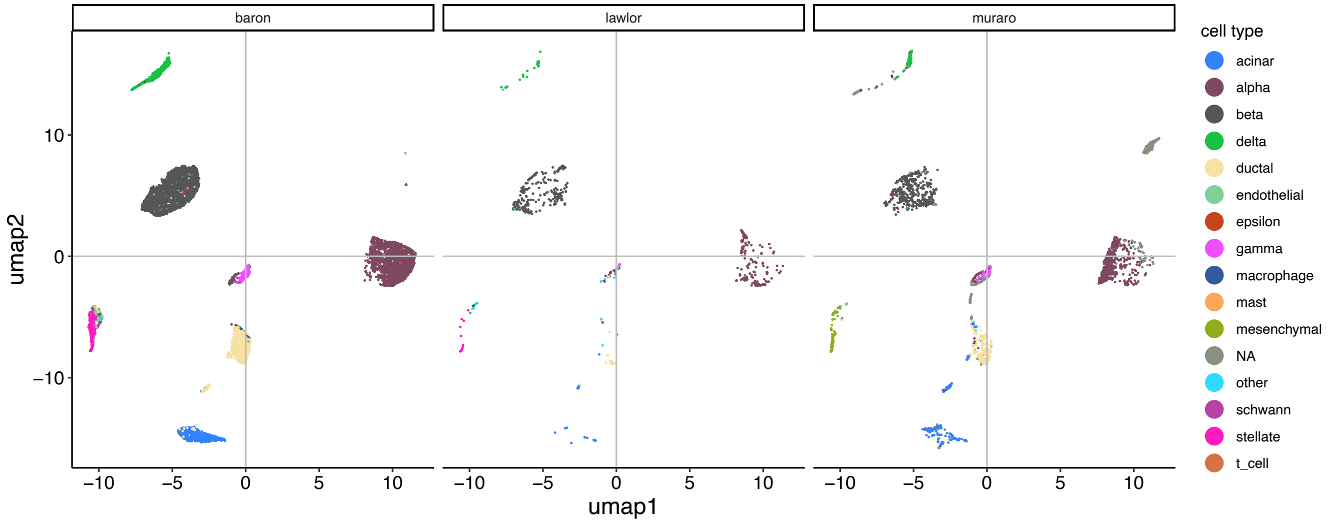

#### D. Seurat: SCTransform with CCA integration

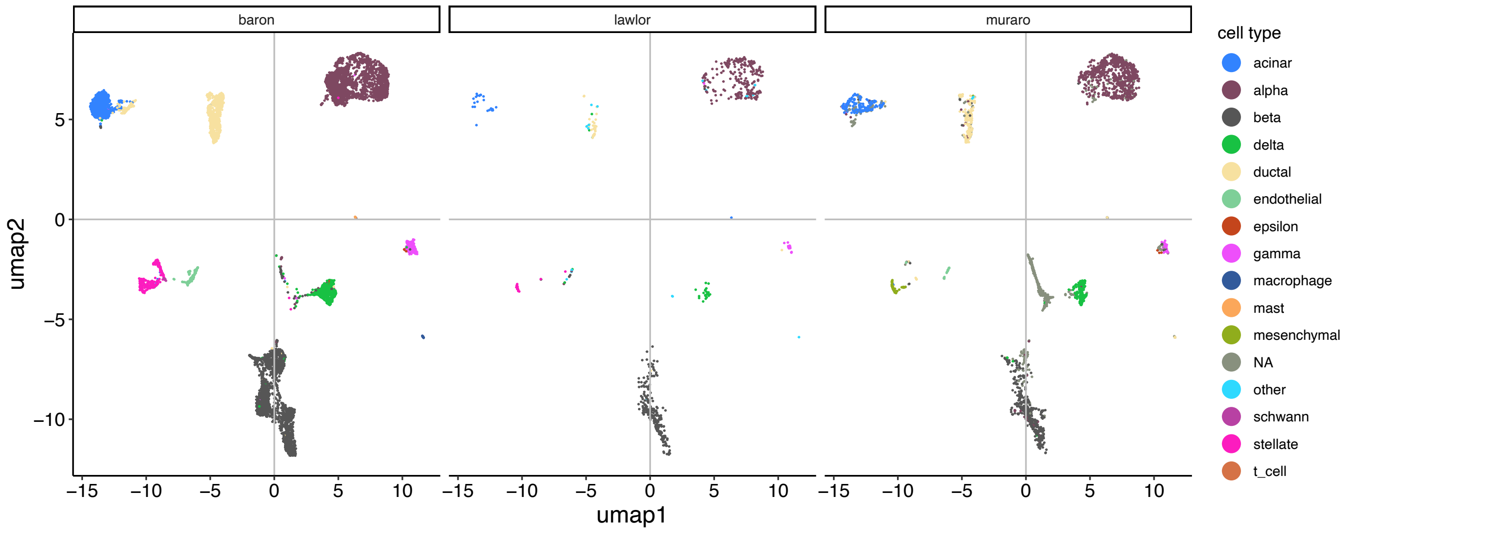


#### E. LIGER

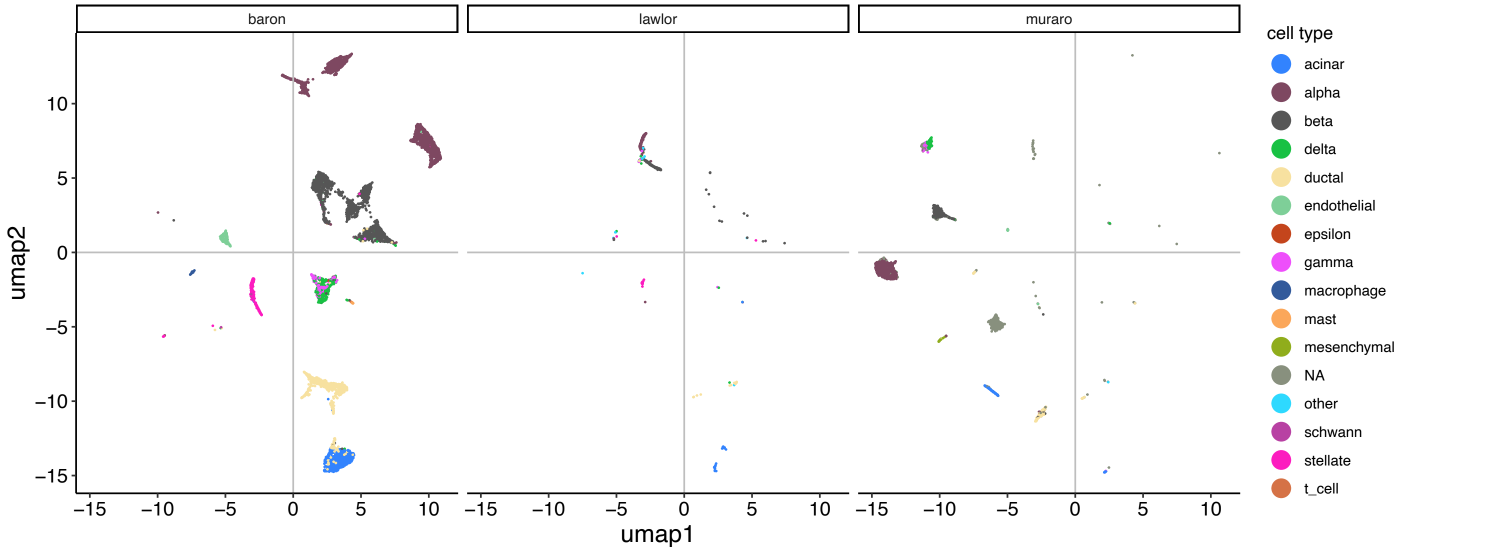


#### F. fastMNN

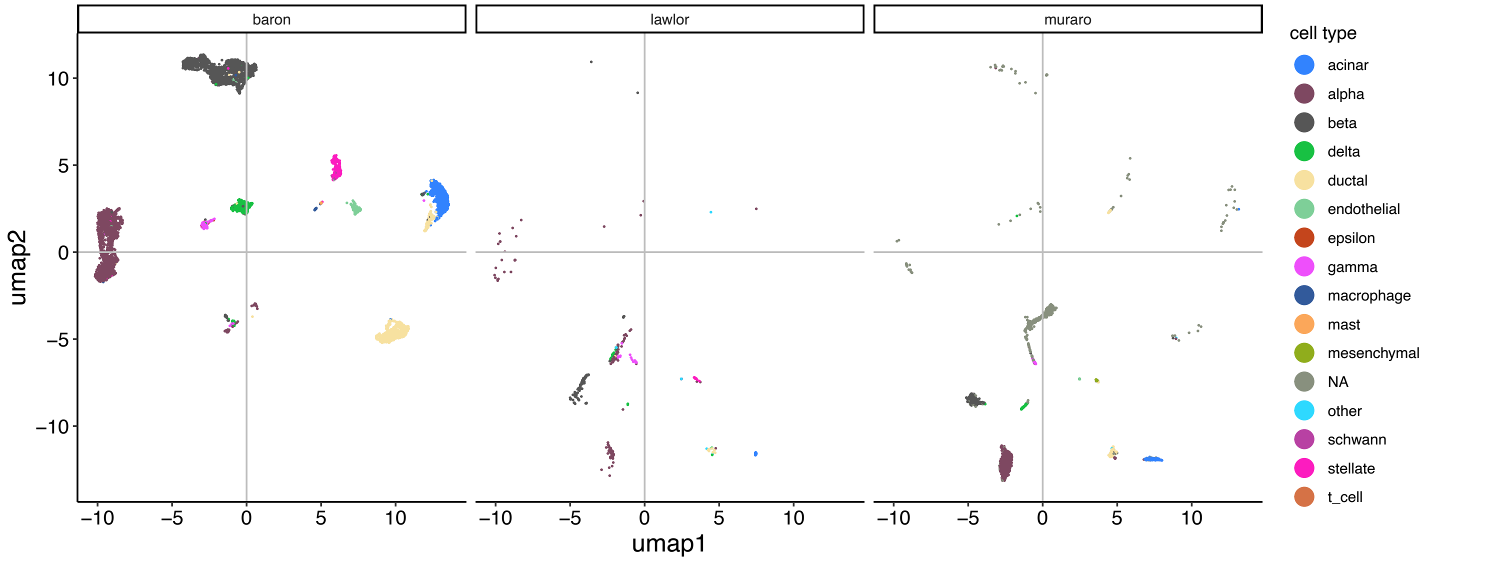

#### G. corralm (Freeman-Tukey residuals) with MNN correction (reducedMNN)

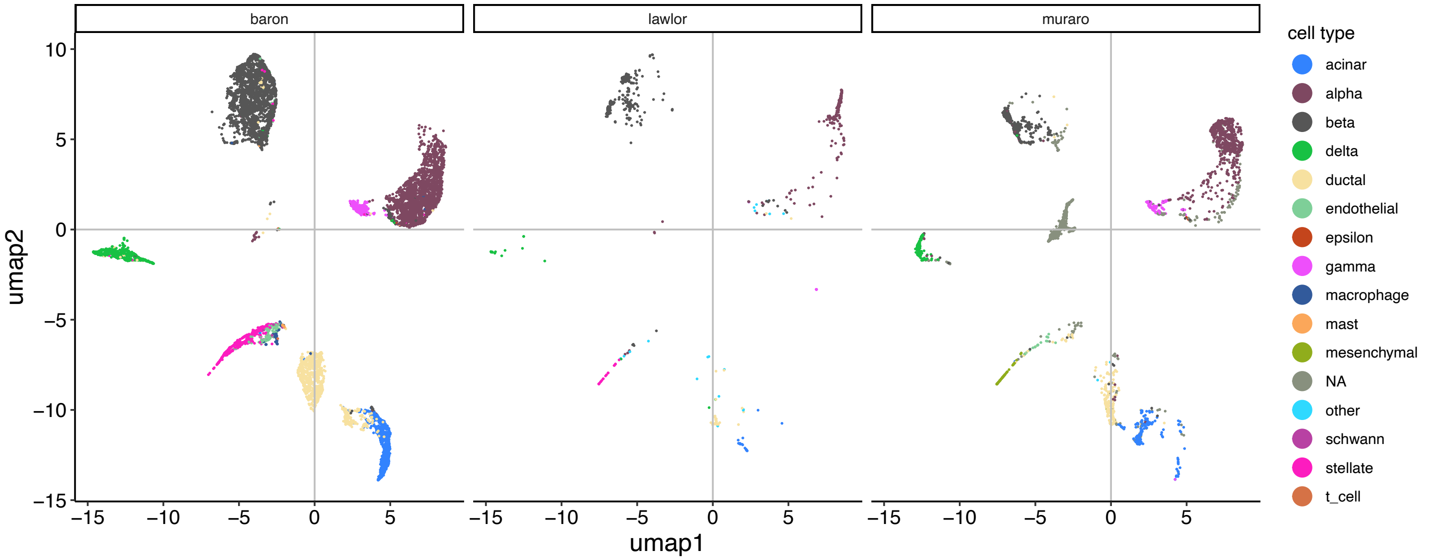


#### H. multibatch PCA

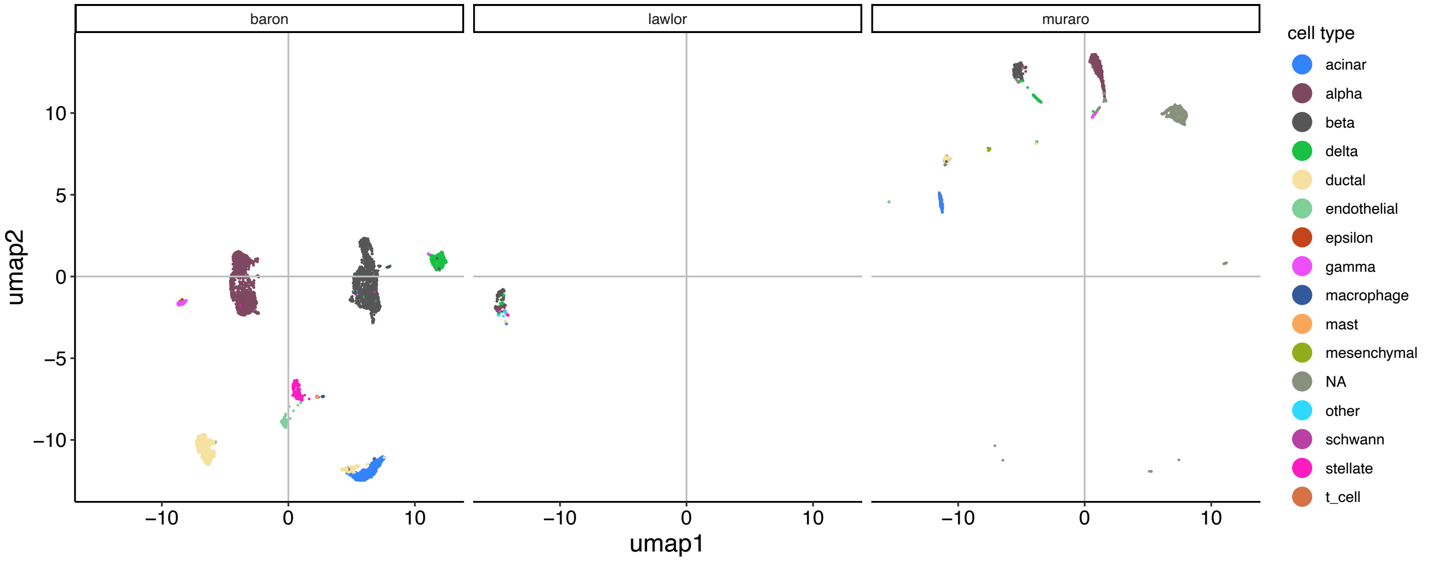


**Figure S5. Pancreas UMAPs, colored by cell type and faceted by dataset batch**. A. corralm on counts; B. Harmony with PCA; C. Harmony with corralm (counts); D. Seurat SCTransform with CCA integration; E. LIGER; F. fastMNN; G. corralm -> reducedMNN; H. multibatch PCA
